# Molecular architecture of OXGR1 reveals evolutionary conserved mechanisms for metabolite surveillance

**DOI:** 10.1101/2025.09.17.676713

**Authors:** Xinyue Zhang, Yujie Lu, Xinheng He, Shimeng Guo, Changyao Li, Yu Wang, Yuan Gao, Juxia Yao, Qingning Yuan, Yinshan Tang, Wen Hu, Kai Wu, Yue Wang, Wanchao Yin, Xin Xie, H. Eric Xu, Heng Liu

## Abstract

The ability of cells to sense and respond to metabolic signals is fundamental to life, yet the molecular mechanisms underlying metabolite surveillance remain incompletely understood. Here, we elucidate the structural basis of metabolite recognition by OXGR1, a GPCR that monitors key intermediates in the tricarboxylic acid cycle (TCA). Using cryo-electron microscopy, we determined four cryo-EM structures of OXGR1 bound to α-ketoglutarate (AKG), itaconate (ITA), and structural related succinate (SUC) and maleate (MA). These structures reveal a positively charged binding pocket and an extensive hydrogen-bond network critical for OXGR1 recognizing dicarboxylic acids. In addition, we identify a distinct arrangement of hydrophobic residues that modulates ligand potency and selectivity. Mutational analysis and molecular dynamics simulations further demonstrate that non-canonical micro-switch motifs, including FRY and NLxxY, are essential for ligand recognition and receptor activation. Comparative structural and evolutionary analyses indicate that these mechanisms are conserved across species, underscoring the critical role of OXGR1 in maintaining metabolic homeostasis. Together, our findings provide a mechanistic framework for metabolite sensing via OXGR1 and suggest potential strategies for therapeutic modulation of metabolic and inflammatory diseases.

## Introduction

Life depends fundamentally on the cell’s ability to monitor and respond to metabolic signals, yet our understanding of how cells achieve this surveillance has remained limited. At the heart of cellular metabolism lies the tricarboxylic acid (TCA) cycle^1^. Despite millions of years of evolutionary changesacross species, the TCA cycle remains the core pathway of cellular energy metabolism^2^. TCA cycle not only generates energy but also produces key metabolites that function as crucial signaling molecules^3^. Among these, α-ketoglutarate (AKG) and itaconate (ITA) have emerged as key regulators that coordinate cellular metabolism with broader physiological responses^4–7^.

The discovery that AKG functions beyond its metabolic role has expanded our understanding of cellular regulation^8,9^. As a required cofactor for over 60 dioxygenases, AKG orchestrates fundamental processes including epigenetic modifications, protein hydroxylation, and fatty acid metabolism^10,11^. Recent studies have revealed AKG’s unexpected role in aging and longevity through regulation of mTOR signaling and mitochondrial function^6,7,12^. This positions AKG at a crucial nexus between metabolism and cellular fate decisions, raising fundamental questions about how cells sense and respond to its levels.

Similarly, ITA as a key immune-metabolite links metabolites to immunity, representing a significant advancement in the field of immune-metabolism^5^. Originally identified as an antimicrobial compound^13^, ITA is now recognized as a key regulator of immune responses^14^. Through inhibition of succinate dehydrogenase and activation of antioxidant pathways via Nrf2 signaling, ITA creates a metabolic environment that suppresses inflammation^4,5,15^. This connection between metabolism and immunity through ITA exemplifies how metabolic signals can reprogram cellular function^16^, but the molecular mechanisms enabling this communication have remained elusive.

OXGR1, residing within the metabolite-sensing δ-branch of class AGPCRs, alongside SUCR1, HCARs, FFARs, and other metabolic sensors (Fig. 1a), has emerged as a crucial node in metabolic recognition^17^. The identification of OXGR1 as a sensor for both AKG and ITA has provided important insights into metabolite surveillance^18,19^. OXGR1 demonstrates that cells could directly monitor TCA cycle intermediates through receptor-mediated signaling^18,20^. The physiological relevance of this surveillance system has become increasingly apparent, with OXGR1 signaling now implicated in diverse processes, including cardiac function^21,22^, adipose tissue metabolism^23^, renal acid-base balance maintaining^24–26^ and inflammatory responses^19,27^. Yet, the fundamental question remained: how does OXGR1 achieve such recognition of structurally distinct metabolites while maintaining the sensitivity required for further cellular responses?

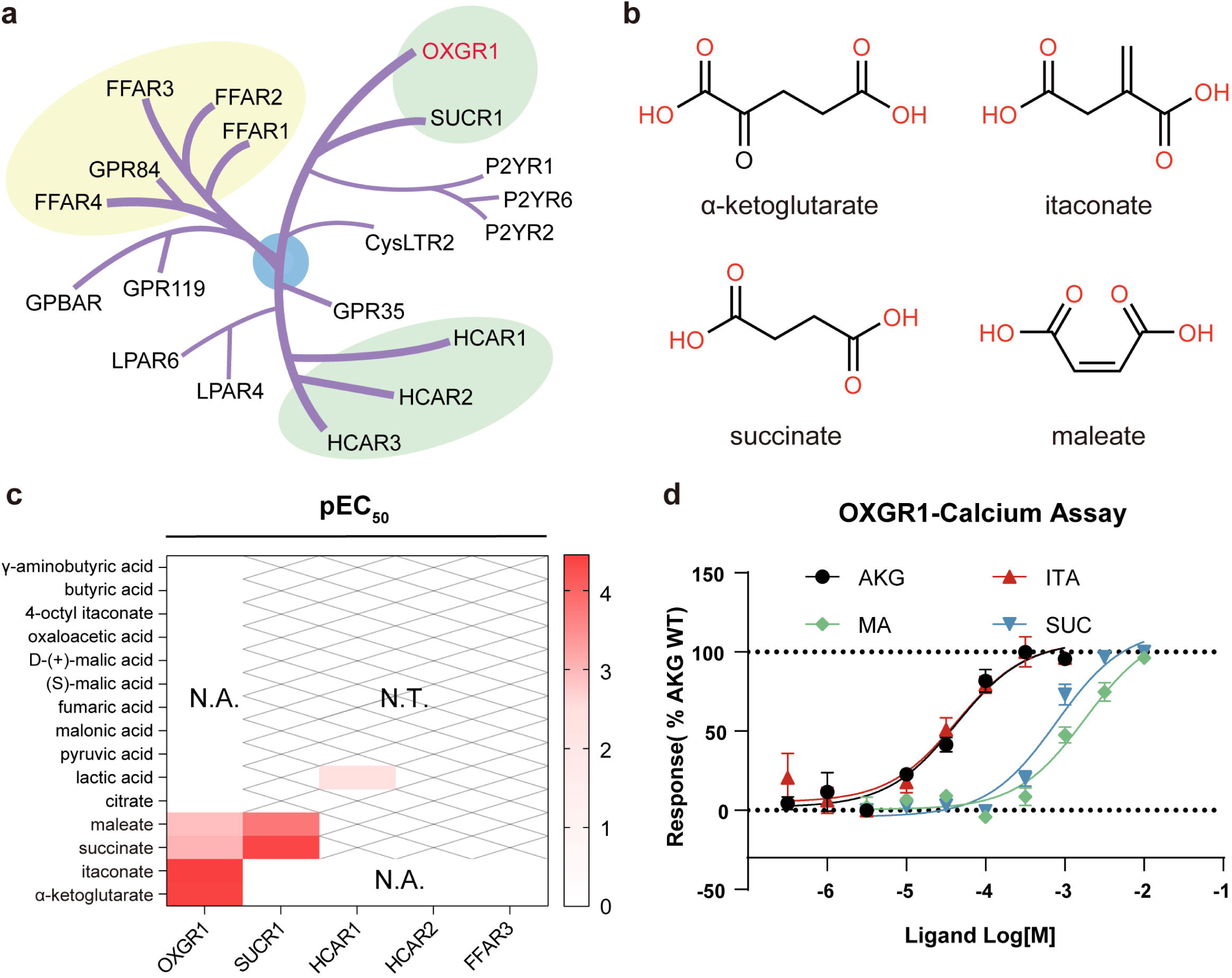

Our work provides molecular insights into metabolite surveillance by resolving four cryo-EM structures of OXGR1 bound to AKG, ITA, as well as the TCA cycle metabolites, succinate (SUC) and maleate (MA), relatively. Through structural analysis combined with functional studies and molecular dynamic simulations, we reveal a specialized binding pocket architecture that enables discrimination between structurally similar metabolites and identify a non-canonical micro-switch in OXGR1 that contribute to receptor activation and facilitate cellular responses to TCA cycle intermediates. These discoveries establish OXGR1 as the prototype of a novel class of metabolite sensors and provide a new framework for understanding cellular surveillance of metabolism. In addition, the structural insights provided a molecular basis for drug development targeting OXGR1 in metabolic and inflammatory diseases.

## Results

### Metabolic surveillance by OXGR1

To comprehensively map OXGR1’s metabolite recognition capabilities, we embarked on a systematic exploration of its activation by TCA cycle intermediates and related compounds. Within 15 metabolically relevant molecules in our experiment (Extended Data Fig. 1a), we find that beyond its known activators AKG and ITA, OXGR1 also responds to succinate (SUC) and maleate (MA) (Fig. 1b, c and Extended Data Fig. 1b). AKG and ITA emerged as the most potent activators, triggering robust G_q_-mediated calcium signaling with EC_50_ values of 49 µM and 43µM respectively, with no detectable activation of G_i_ or G_s_ signaling (Fig. 1c, d and Extended Data Fig. 1b, c). SUC and MA also activated OXGR1, albeit with lower potency, EC_50_ values of 800 µM and 2000 µM, respectively, exclusively through the G_q_ pathway (Fig. 1c and Extended Data Fig. 1b). This selective coupling to G_q_ suggests a specialized signaling mechanism tuned to metabolic surveillance.

Under normal physiological conditions, the plasma concentrations of AKG in mice typically range from 50 to 70 μM. Physiological or metabolic stress, such as physical exercise, can lead to elevated levels of AKG, reaching 90-110 μM^23^. ITA is present at very low concentrations in a resting state, making it difficult to detect. However, during inflammatory responses, particularly in activated macrophages (in mouse bone), the concentration of ITA can significantly increase, even reaching 5 mM^28^. The concentrations of succinate in blood are generally around 6 to 32 μM (HMDB: HMDB0000254). In pathological conditions such as obesity, hypoxia, inflammation, or metabolic disturbances, succinate levels can rise to between 150 and 300 μM^29–31^. MA is typically found at low concentrations in the body. These studies, combined with our results, indicate that under physiological conditions, OXGR1 primarily senses AKG and ITA.

Further, we examined the ligand’s activation profile against related carboxylic acid-sensing GPCRs, including SUCR1, HACR1/2, and FFAR3 (Fig. 1c and Extended Data Fig. 1d). Strikingly, while SUC and MA showed expected activation of SUCR1, AKG displayed no cross-reactivity, and ITA exhibited only minimal activation of SUCR1 and no detectable activation of other related receptors (Fig. 1c and Extended Data Fig. 1d).

Overall, OXGR1 exhibits relatively high activity in response to AKG and ITA, while showing markedly lower response to structurally related metabolites, SUC and MA, along with selective coupling to G_q_-mediated signaling. Notably, both AKG and ITA are specifically recognized by OXGR1. These findings reveal a molecular mechanism by which OXGR1 discriminates among closely related metabolites, enabling selective recognition of specific metabolites within the complex cellular milieu.

### Architecture of OXGR1 for metabolite capture and recognition

To unravel the molecular mechanisms underlying OXGR1’s distinct metabolite discrimination, we determined four cryo-EM structures of OXGR1 bound to AKG, ITA, SUC, and MA. Through protein engineering and complex assembly, we successfully obtained stable OXGR1-G_q_ complexes in their active states. We employed a BRIL fusion strategy at the receptor’s N-terminus and assembled the complex with engineered Gα_q_, rat Gβ_1_, and bovine Gγ_2_, further stabilized by scFv16 and Nb35 antibody fragments ^32^(Extended Data Figs. 2, 3).

We determined cryo-EM structures of OXGR1-G_q_ complexes, bound with AKG, ITA, SUC, and MA, at overall resolutions of 2.66 Å, 2.65 Å, 2.64 Å, and 2.73 Å, respectively. To improve the visualization of ligand densities, we performed focused refinement on the receptor, yielding local resolutions of 2.89 Å, 2.90 Å, 2.70 Å, and 2.97 Å for the respective OXGR1-ligand complexes (Fig. 2a-d, Extended Data Figs. 2-4 and Supplementary Table 1, 2). These high-resolution maps clearly show most portion of OXGR1, the G_q_ heterotrimer, stabilizing antibody fragments, and the respective ligands (Fig. 2a-d). In all four structures, the ligand densities are visible and of sufficient quality to allow modeling of the bound metabolites, but the precise orientation of certain ligands remains uncertain. To further validate the ligand binding poses, we compared our experimental structures with molecular docking models of ligand-bound OXGR1, and found that the ligand positions are largely consistent (Extended Data Fig. 5).

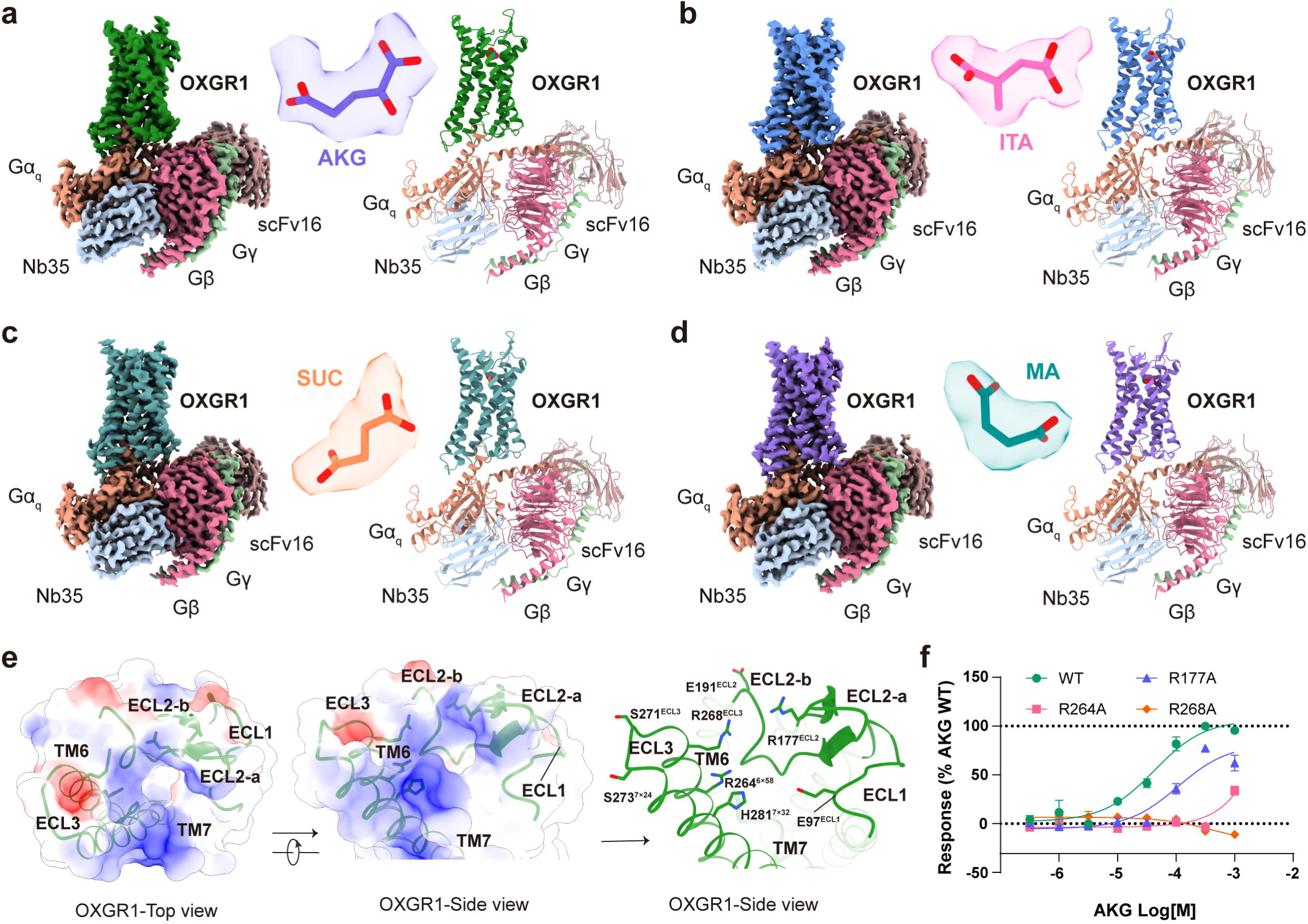

In the OXGR1 structures, the second extracellular loop (ECL2) is structurally divided into two parts: ECL2-a (I172^4x64^ - L184^45x51^) forms a β-hairpin over the binding pocket, while ECL2-b (D185^45x52^ - E191^ECL2^) loop integrates into and contributes to formation of the ligand-binding pocket (Fig. 2e). This arrangement also observed in related receptors like SUCR1^33–35^ and lipid sensors, such as DP2^36^ and BLT1^37^ (Extended Data Fig. 6a).

Notably, the extracellular surfaces of TM6, TM7, and ECL2-a create a positively charged vestibule through key residues H281^7x32^, R264^6x58^, R268^6x62^, and R177^ECL2^ (Fig. 2e). The functional importance of this charge distribution was highlighted by mutational analysis, where alanine substitutions of the positively charged residues significantly reduced or entirely eliminated receptor activation (Fig. 2f). The spatial distribution and structural features of these basic amino acids may be associated with ligand recognition and entry in OXGR1, similar to the reported “cationic lure” entry mechanisms observed in lipid receptors, such as GPR84 and BLT1^17,37,38^(Extended Data Fig. 6b).

### Dicarboxylic metabolites recognized by OXGR1

The orthosteric binding pocket of OXGR1 is composed by TM1, TM2, TM3, TM7 and the extracellular loop ECL2 (Fig. 3a and Extended Data Fig. 6c). Notably, the pocket exhibits a unique dual-nature architecture, with an electropositive upper region that transitions into a hydrophobic base (Fig. 3a and Extended Data Fig. 6c), creating an optimal environment for capturing and orienting dicarboxylic acid metabolites. To better describe the interactions between ligands and receptor, OXGR1 binding pocket is divided into three sub-pockets: SP1 adjacent to TM3, SP2 near TM1/TM7, and SP3 forming a hydrophobic floor (Fig. 3a and Extended Data Fig. 6c). This tripartite arrangement proves crucial for ligand discrimination, with each sub-pocket playing an important role in metabolite recognition and orientation. At the top of the ligand-binding pocket, a polar hydrogen-bond network, involving D185^45x52^, Y93^2x54^, H281^7x32^, and E97^ECL1^, is mediated by two water molecules, W1 and W2, and extends toward the extracellular space (Fig. 3b and Extended Data Fig. 6d). Mutational analyses that disrupt this hydrogen-bond network result in weakened or completely abolished ligand-induced receptor activation, highlighting its critical role in ligand recognition and receptor activation (Fig. 3c).

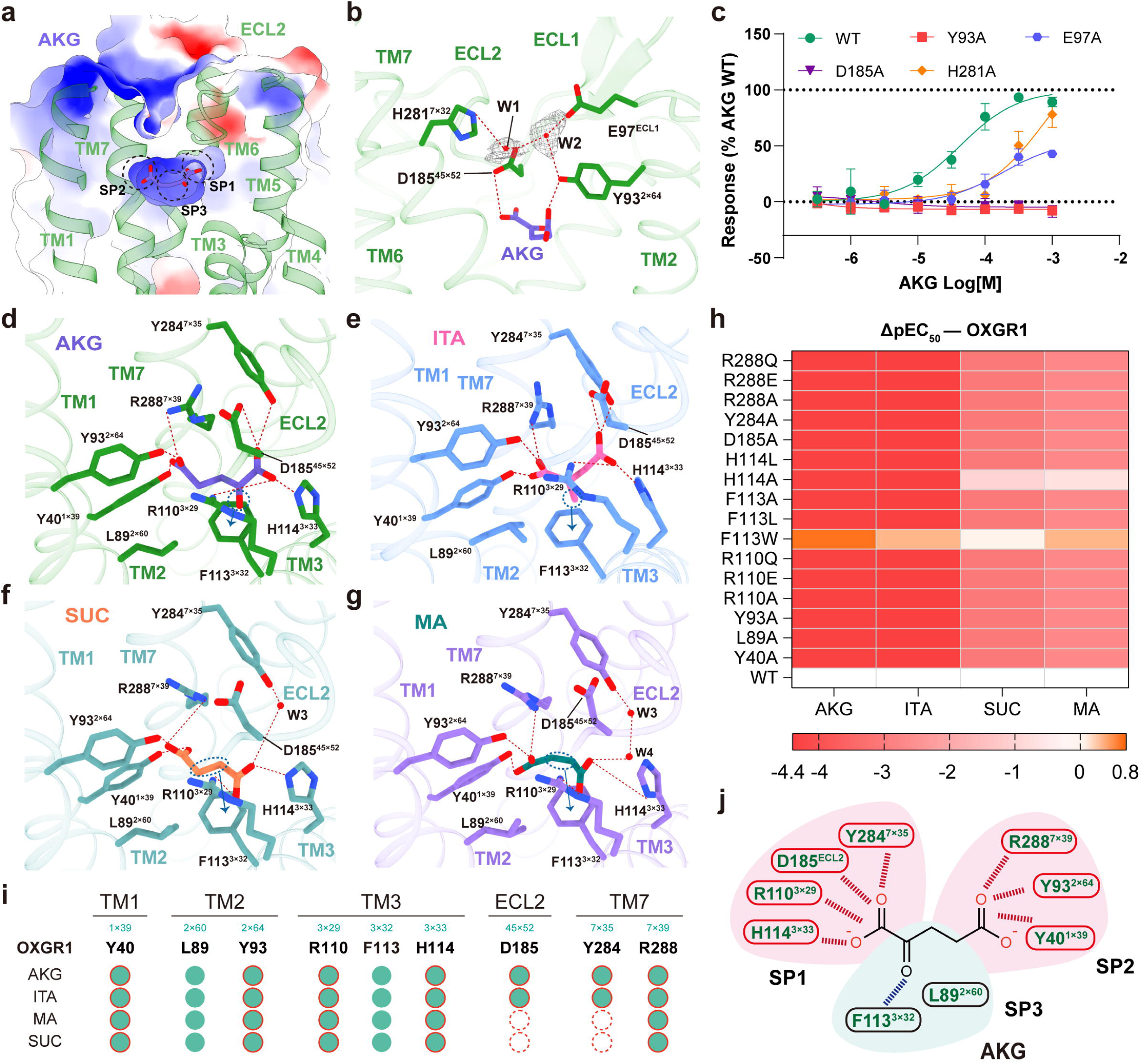

In the binding pocket, all four metabolites, including AKG, ITA, SUC, and MA, adopt a conserved binding mode, characterized by the insertion of their terminal carboxyl groups into sub-pockets SP1 and SP2, where they form an extensive network of polar interactions. The central linker region of each ligand extends into SP3, engaging in additional hydrophobic or π-stacking interactions (Fig. 3a and 3d-g). In the AKG and ITA bound structures, the R110^3x29^, H114^3x33^, D185^45x52^ and Y284^7x35^ coordinate one carboxyl group in SP1, while R288^7x39^, Y40^1x39^, and Y93^2x64^ anchor the second carboxyl group in SP2 (Fig. 3d-g and Extended Data Fig. 6e). In the SUC-OXGR1 complex, interactions between SUC and Y284^7x35^ and D185^45x52^ are connected by one water molecule (W3); whereas in the MA-OXGR1 complex, two water molecules (W3 and W4) mediate these interactions. At the bottom of the binding pocket, the connected alkyl chains in four ligands, varying in length and functional groups, interact with F113^3x32^ and L89^2x60^ (Fig. 3d-g). The functional relevance of this recognition system was supported by comprehensive mutational analysis. Notably, nearly all mutations of binding pocket composing residues completely abolished receptor activation, while maintaining normal expression levels (Fig. 3h and Extended Data Fig. 7; Supplementary Table 3). This may result from the inherently low activity of the ligand-activated receptor, rendering partial loss of function difficult to detect, or disruption of molecular coordination due to the mutations.

Through systematic comparison of four ligand-bound structures(Fig. 3i), we identified the core recognition machinery for dicarboxylic metabolites recognition: residues R288^7x39^, Y93^2x64^, Y40^1x39^, R110^3x29^ and H114^3x33^ form the primary polar interaction network with the ligands’ carboxylate groups (Fig. 3f-g). In addition, the central linker region of the ligands engages in hydrophobic interactions with F113^3x32^, further stabilizing ligand binding (Fig. 3j and Extended Data Fig. 6e). The proposed charge-charge interactions promote deprotonation of the carboxyl group on AKG, as confirmed by Epik pKa calculations^39^. This configuration provides a structural basis for binding of dicarboxylic metabolites to receptor-activating conformational changes. The recognition of AKG by OXGR1 was previously modeled based on the HCAR2 receptor^40^, in which AKG was docked in a vertical orientation, resembling the pose of nicotinic acid in HCAR2. In contrast, our experimentally determined cryo-EM structures reveal that AKG adopts a horizontal binding mode, similar to that of succinate in SUCR1. In this configuration, the two carboxylate groups insert into distinct sub-pockets (SP1 and SP2), while the central linker engages SP3 through hydrophobic and water-mediated interactions.

Structural comparisons among OXGR1, SUCR1, and HCAR2/3 further reveal a clear distinction in ligand binding orientations^34,41,42^. Dicarboxylic acid receptors such as OXGR1 and SUCR1 accommodate ligands in a parallel-to-membrane orientation, enabling dual-site anchoring. In contrast, monocarboxylic acid receptors like HCAR2/3 bind ligands such as nicotinic acid perpendicular to the membrane, inserted into the orthosteric pocket (Fig. 5c and Extended Data Fig. 8). These contrasting orientations reflect fundamentally different recognition strategies shaped by ligand chemistry and receptor architecture.

### Mechanisms underlying differential ligand potency

The marked difference in OXGR1 activation potency, approximately 20-fold higher for ITA and AKG compared to MAand SUC (Fig. 1c, d and Extended Data Fig. 1b), prompted a detailed investigation into the molecular determinants underlying this disparity. Using integrated computational and experimental approaches, we established a binding energy hierarchy for OXGR1 ligands (ITA >AKG > SUC≈MA), based on six 1,000-ns molecular dynamics (MD) simulations (Fig. 4a). ITA exhibited the highest binding stability, with a binding free energy of -52.1 ± 0.9 kcal/mol (Fig. 4a), followed by AKG (-40.3 ± 3.3 kcal/mol). In contrast, SUC and MA displayed substantially weaker binding affinities (-31.8 ± 2.5 kcal/mol and -29.7 ± 6.0 kcal/mol, respectively) (Fig. 4a). This dynamic behavior provides a molecular basis for the reduced potency of SUC and MA. Further structural and predicted ranking of binding free energies revealed that residues F113^3x32^, D185^45x52^, and Y284^7x35^ contribute differentially to ligand binding, depending on the specific chemical features of each metabolite (Fig. 4b-i).

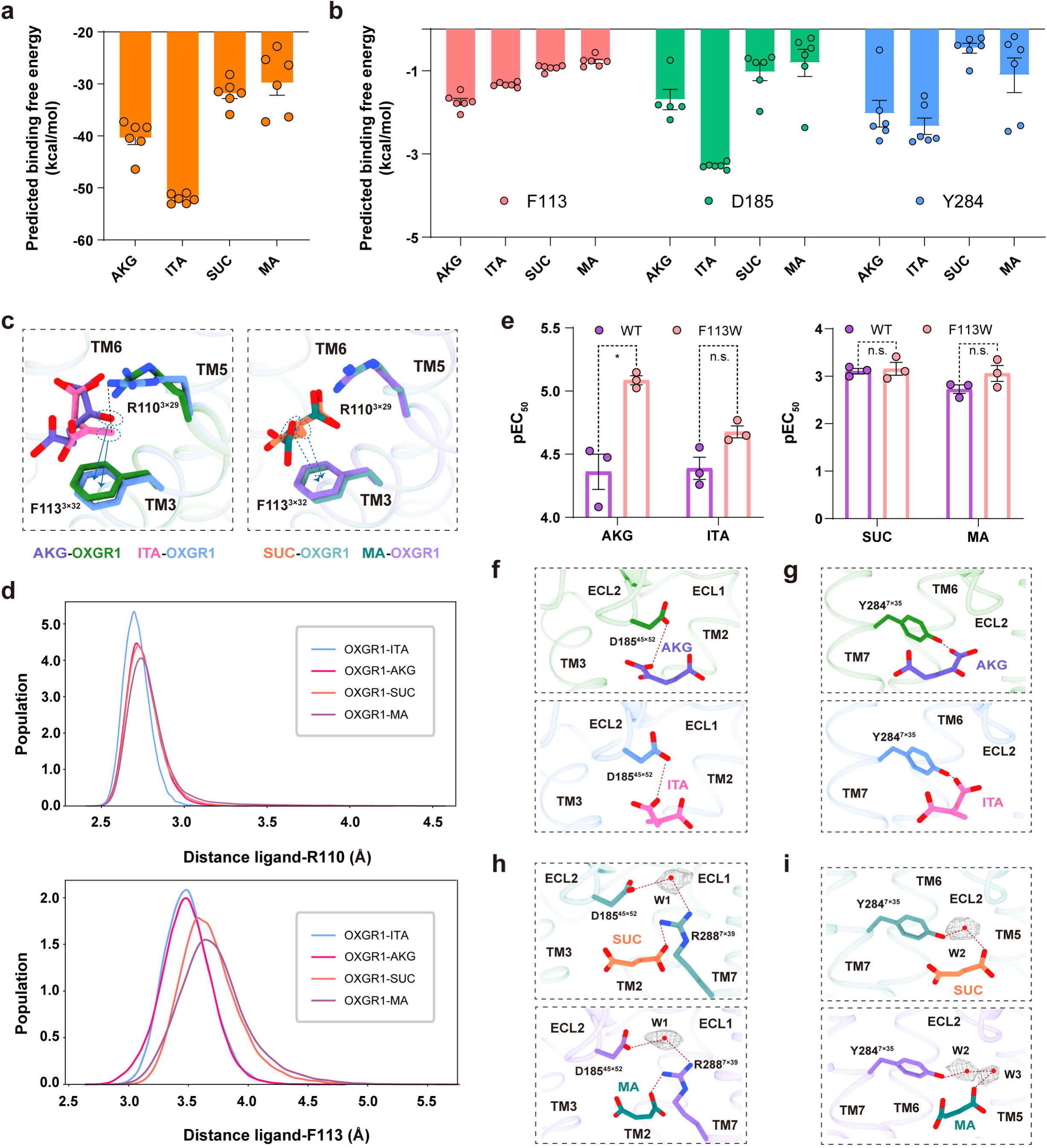

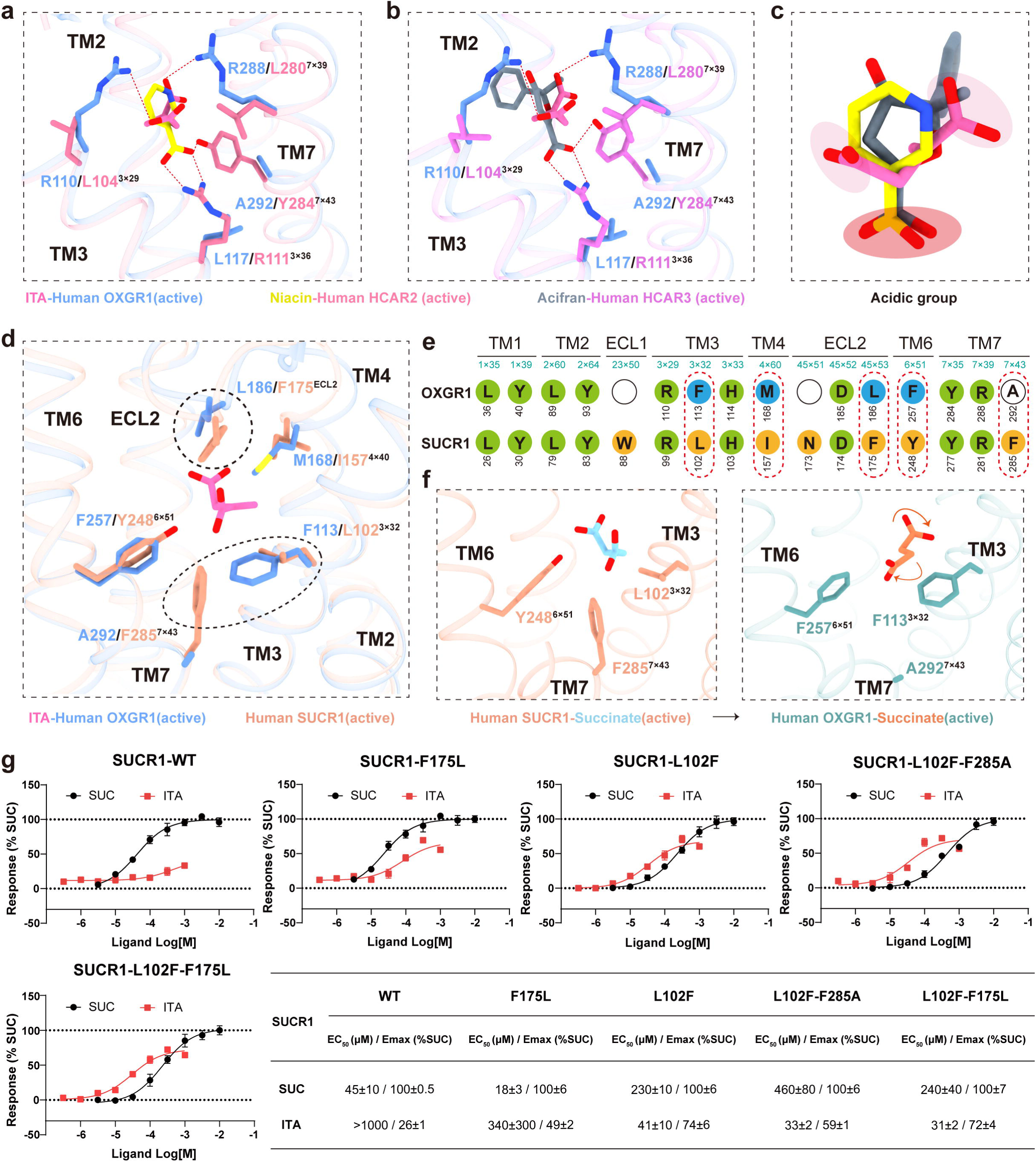

The ketocarbonyl group in AKG and the double bond in ITA represent key structural features that underlie their enhanced potency. In the AKG-bound structure, the ketocarbonyl group engages in a σ-π interaction with F113^3x32^, contributing to pocket stabilization (Fig. 4c-e). In the case of ITA, the presence of a conjugated double bond between R110^3x29^ and F113^3x32^ generates a distinct electronic environment that facilitates electrostatic interactions with R110^3x29^ and π-π stacking with F113^3x32^(Fig. 4c-e). Consistent with these structural observations, molecular dynamics simulations showed that interactions between F113^3x32^, R110^3x29^ and the ligands were stably maintained throughout the trajectories (Fig. 4d). To functionally validate the importance of these interactions, we introduced a F113W mutation to enhance the aromatic and electronic interactions within the pocket. Functional assays revealed that this mutation led to a ∼6-fold increase in AKG potency and a ∼2-fold increase for ITA. Notably, MA, due to the presence of a double bond, also exhibited ∼2-fold enhanced potency, while SUC, lacking these features, showed no significant change (Fig. 4e).

Additionally, these features increase AKG’s and ITA’s rigidity, and expand the interaction interface, enabling direct hydrogen bonding with residues D185^45x52^ and Y284^7x35^(Fig. 4f, g). In contrast, SUC and MA lack such structural elements and exhibit smaller molecular volumes, resulting in weaker interactions with these key residues that are instead mediated by structured water molecules (Fig. 4h, i). These findings are supported by molecular dynamics simulations, which predict stronger binding free energies for AKG and ITA compared to SUC and MA (Fig. 4b). Collectively, these results underscore the critical roles of F113^3x32^, R110^3x29^, D185^45x52^ and Y284^7x35^ in mediating structure-based discrimination among dicarboxylic metabolites.

### Structural determinants of ITA selectivity

The pronounced selectivity of ITA and AKG for OXGR1 over related metabolite-sensing receptors presents a valuable case study in molecular recognition and selectivity. In our studies, comparative analysis of binding pocket architectures between OXGR1 and related HCAR structures revealed significant differences in ligand orientation and pocket composition. While HCAR receptors orient their ligands vertically, with the carboxylate anchored at the base, OXGR1 uniquely positions ITA’s dual carboxyl groups horizontally across the pocket (Fig. 5a-c). This distinct binding mode is enabled by OXGR1’s enriched basic residue environment, including R110^3x29^, R288^7x39^, and H114^3x33^, contrasting sharply with the single arginine found in HCAR2/3 pockets (Extended Data Fig. 9).

Notably, comparison with SUCR1, another receptor involved in dicarboxylic acid sensing, revealed key structural determinants underlying ITA selectivity, particularly in differentiating responses to ITA. Comparison of OXGR1 and SUCR1 structures reveal that the arrangement of key polar residues (R^7x39^, Y^2x64^, Y^1x39^, R^3x29^, and H^3x33^) responsible for dicarboxylic acid recognition is highly conserved within the binding pockets of OXGR1 and SUCR1 (Fig. 5d, e and Extended Data Fig. 9), indicating a shared structural basis for metabolite recognition. In contrast, the surrounding hydrophobic environment differs markedly between the two receptors. OXGR1 contains M168^4x60^, L186^ECL2^, F257^6x51^, A292^7x43^, and F113^3x32^, whereas the corresponding positions in SUCR1 are occupied by I157^4x60^, F175 ^ECL2^, Y248^6x51^, F285^7x43^, and L102^3x32^ (Fig. 5d, e). This architectural difference allows OXGR1 to accommodate ITA’s rigid and extended backbone, facilitating favorable electrostatic and π-stacking interactions within the binding site. In contrast, the tighter pocket of SUCR1 imposes steric constraints that restrict the binding pose of larger or more rigid ligands such as ITA (Fig. 5f).

To further evaluate the role of the hydrophobic pocket in governing ITA selectivity, we performed structure-guided engineering of SUCR1. By introducing key hydrophobic residues from OXGR1 into SUCR1, we generated a series of gain-of-function mutants. Of note, the SUCR1-F175^ECL2^L mutation enabled partial ITA responsiveness, while SUCR1 -L102^3x32^F, SUCR1-L102^3x32^F-F285^7x43^A as well as SUCR1-L102^3x32^F-F175^ECL2^L mutations further enhanced ITA activation in both potency and efficiency (Fig. 5g, Extended Data Fig. 10 and Supplementary Table 4). These results functionally validate the structural observations, demonstrating that the specific arrangement and size of hydrophobic residues are key determinants for shaping the binding pocket geometry required for effective ITA recognition.

### Unique micro-switch motifs for OXGR1 activation

All four ligand-bound OXGR1 structures are in complex with G proteins. Compared to the antagonist-bound SUCR1 structures, OXGR1, as exemplified by the AKG bound complex, adopts a fully active conformation as characterized by the pronounced outward displacement of TM6 at the cytoplasmic end and rotation of TM5 (Extended Data Fig. 11a), hallmark features of GPCR activation that create the G protein binding interface^43^.

At the molecular level, detailed analysis revealed that OXGR1 employs non-canonical GPCR micro-switch motifs to mediate its activation. While most class A GPCRs (86%) employ a D(E)RY motif to form an ionic lock stabilizing the inactive state^44–46^ (Fig. 6a), OXGR1 features a non-canonical FRY motif. This substitution eliminates the ionic lock, allowing F130^3x49^ to engage a hydrophobic pocket formed by I71^2x42^, F127^3x46^, and I144^34x56^ (Extended Data Fig. 11b). Functional assays revealed that mutating F130^3x49^ to D/E abolished activation by all ligands (Fig. 6b), supporting the importancerole of this motif in activation. Molecular dynamics simulations further illuminated how this unique micro-switch architecture shapes receptor behavior. During simulations, WT-OXGR1 exhibits a greater outward displacement of TM6 compared to F130^3x49^D-OXGR1, indicating the FRYmotif promotes active stateconformations. In contrast, the F130^3x49^D substitution disrupted thelocal hydrophobic network, increasing side-chain mobility and causing TM6 retraction towards intermediate state (Fig. 6c). A second unique featureappears in the NPxxYmotif, where OXGR1 replaces the highly conserved (96%) proline with leucine at position 7x50^44,45^ (Fig. 6a). This substitution reduces TM7 kinking compared to other class A GPCRs (Extended Data Fig. 11c), enabling a 2.2 Å greater inward movement of TM7 (Extended Data Fig. 11a).

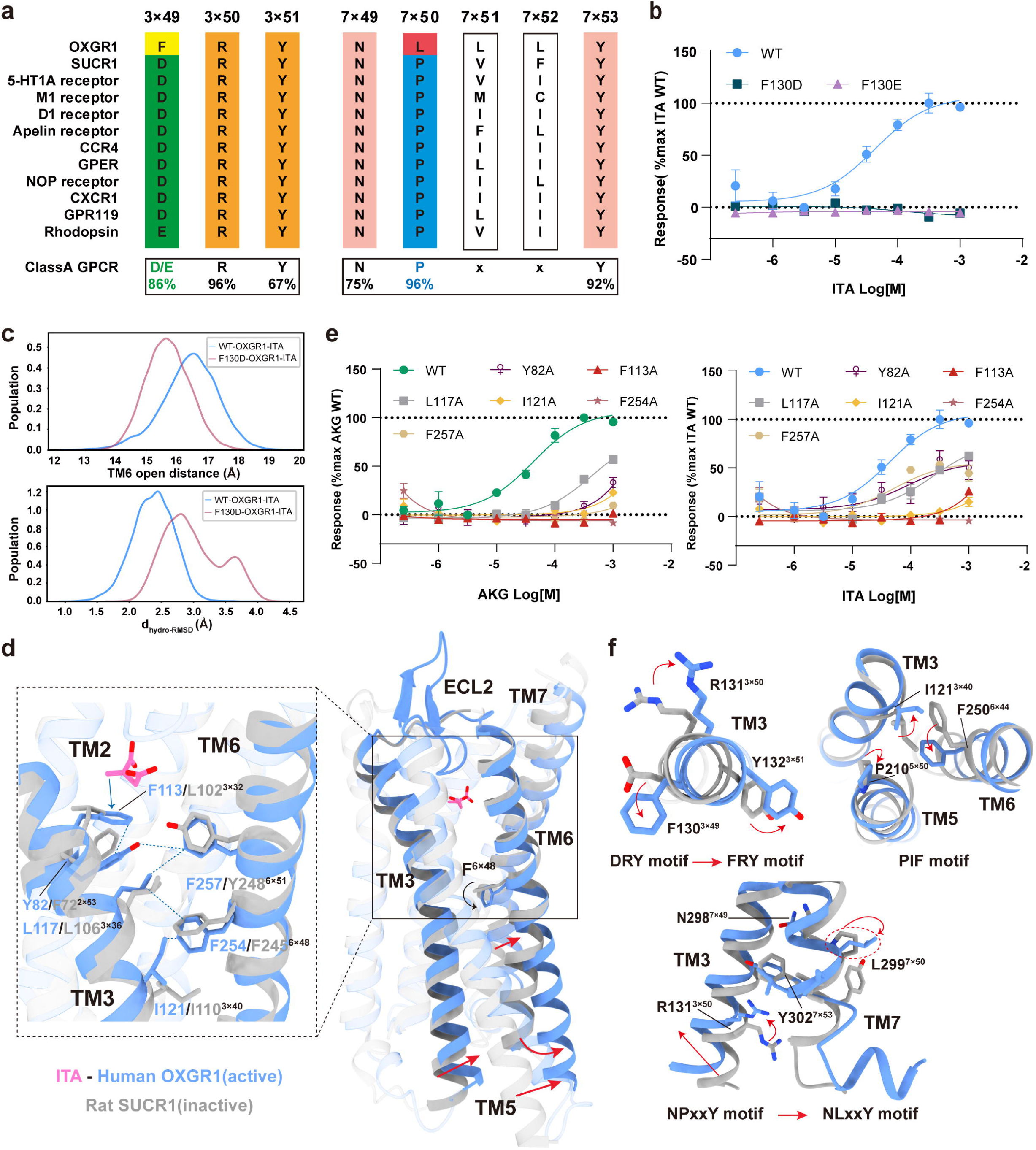

In contrast to the HCAR family^41,42^, in which ligand binding primarily leads to conformational changes affecting residues such as R^3x36^ and Y^7x43^, ligand binding in OXGR1 involves direct interaction between AKG and F113^3x32^. This initial contact triggers signal propagation through a hydrophobic network including Y82^2x53^, F257^6x51^, I121^3x40^, and L117^3x36^, ultimately reaching the F254^6x48^ toggle switch (Fig. 6d). Functional assays confirmed that mutations in these key signaling residues variably impaired receptor activation, emphasizing their essential role in transducing the activation signal (Fig. 6e). The rearrangement of F^6x48^ is accompanied by a marked displacement of F257^6x44^ toward TM5 in the PIF motif (Fig. 6f) and further associated with the displacement of the NPxxY motif toward the center of the core. Together, these conformational changes contribute to the outward displacement of TM6, and the adjacent TM5 and TM7, accommodating the G_q_ coupling of OXGR1(Extended Data Fig. 11a).

### Evolutionary insights into OXGR1-mediated metabolic sensing

The evolutionary importance of TCA cycle intermediates^47^, combined with the ability of OXGR1 to selectively sense AKG and ITA under physiological conditions, raises the question of whether its ligand recognition and activation mechanisms are conserved across species. To address this, we performed a comprehensive sequence alignment of OXGR1 orthologs across vertebrates, where the receptor is predominantly expressed.

Sequence analysis revealed a high degree of conservation among residues critical for ligand coordination and receptor activation. Key residues mediating interactions with the dicarboxylic acid moiety, R110^3x29^, R288^7x39^, Y40^1x39^, and D185^45x52^ were almost fully conserved across all examined species, while Y93^2x64^ and H114^3x33^ showed divergence in birds and fish (Extended Data Fig. 12). In addition, F113^3x32^, which stabilizes the ligand linker region and contributes to shaping the binding pocket geometry, was strictly conserved. With regard to activation-related residues, including F113^3x32^, as well as I121^3x40^ and F254^6x48^, was universally conserved, supporting its essential role in propagating ligand-induced conformational changes (Extended Data Fig. 12). Besides, the non-canonical FRY motif (F130^3x49^-R131^3x50^-Y132^3x51^) in OXGR1, which likely facilitates disruption of the inactive-state and promotes G protein coupling, and the NLxxY motif (L299^7x50^), were also conserved. Together, these findings demonstrate that both the ligand recognition architecture and the activation machinery of OXGR1 are evolutionarily conserved, highlighting its central role in sensing dicarboxylic metabolites, particularly ITA and AKG.

## Discussion

AKG and ITA are critical intermediates of the TCA cycle that bridge energy metabolism, cellular signaling, and immune regulation. Both metabolites have been reported to signal through the G protein-coupled receptor OXGR1, influencing processes such as energy balance, acid-base homeostasis, and inflammatory responses. The previous study proposed a homology-based model of OXGR1 recognizing AKG^40^, providing initial insights into the potential binding mode through structural modeling and functional validation. However, the precise molecular mechanisms by which OXGR1 selectively senses AKG and other dicarboxylic acids remained to be experimentally elucidated.

In this study, we present four high-resolution cryo-EM structures of OXGR1 in complex with AKG, ITA, succinate (SUC), and malate (MA), providing a comprehensive structural framework for understanding ligand recognition and receptor activation. We identify a positively charged ligand-binding pocket composed of key polar residues, R110^3x29^, H114^3x33^, Y94^7x35^, Y40^1x39^and R288^7x39^, that form a conserved hydrogen-bonding network essential for anchoring the carboxylate groups of dicarboxylic acids. Leveraging the functional divergence between OXGR1 and its paralog SUCR1, known to sense SUC, but not ITA in our study, we performed structure-guided cross-receptor mutagenesis. This approach uncovered a set of surrounding hydrophobic residues, including F113^3x32^, A292^7x43^, and L186^ECL2^, that not only help maintain the structural integrity of the binding pocket but also fine-tune ligand selectivity and potency by modulating pocket shape and environment.

During the preparation of this manuscript, a cryo-EM structure of OXGR1 bound to AKG (PDB ID: 8YYW) was released. In that structure, two AKG molecules were modeled, one in the orthosteric site and another near the pocket entrance close to ECL2. While the position of AKG in the orthosteric site partially overlaps with our structure, the additional density near ECL2 modeled as AKG in 8YYW corresponds to two well-resolved water molecules in our map (Extended Data Fig. 13) . Thesedifferencesmay reflect the inherent dynamics of ligand binding or arise from differences in local map resolution (2.89 Å in our structure for the receptor region versus 3.16 Å reported in 8YYW). Importantly, our experimental structures capture a conserved ligand-binding mode across four dicarboxylic acids, supported by structure studyand functional mutagenesis, thereby providing a validated and unified structural framework for dicarboxylic acids recognition.

Beyond ligand recognition, our structural analyses reveal that OXGR1 adopts a non-canonical activation mechanism. Unlike most class A GPCRs, which harbor a conserved D(E)RY motif, OXGR1 contains an atypical FRY sequence. This substitution disrupts the classical ionic lock, favoring an active-like conformation characterized by outward displacement of TM6 and enabling G protein coupling. In addition, the replacement of the conserved proline within the NPxxY motif by leucine (L299^7×50^) alters the bending dynamics of TM7, further stabilizing the receptor in an active conformation. Further evolutionary analysis revealed a high degree of conservation in both ligand-binding and activation-related residues across vertebrates, highlighting OXGR1’s evolutionarily preserved role as a dedicated G_q_-coupled sensor for dicarboxylic metabolites.

In conclusion, this study provides critical insights into the structure and function of OXGR1, confirming its evolutionary conservation as a key regulator of metabolic processes. Our findings pave the way for future research exploring OXGR1’s potential as a therapeutic target, particularly in diseases characterized by disrupted metabolic and immune pathways.

### Materials and Methods Constructs

The wild-type human OXGR1 (residues 1-337) was cloned into pFastBac vector with an N-terminal HA signal peptide sequence followed by a Flag tag, a 15×His tag and a thermo-stabilized BRIL to facilitate expression and purification^48^. The Gα_q_ was designed based on a miniGα_s_ skeleton with the N-terminus replaced by Gα_i1_ for binding of the antibody fragments scFv16^49^. All the constructs, including rat Gβ_1_, bovine Gγ_2_ and scFv16, were cloned into pFastBac vector separately.

The full-length human OXGR1 and mutations were fused after HA signal peptide and Flag tag. The constructs were cloned into the pcDNA3.0 vector for the HEK293T system. The full-length human SUCR1 and mutations were fused after HA signal peptide and Flag tag. The constructs were cloned into pcDNA6.0 vector for HEK293T system.

### Cell culture and transfection

HEK293 cells were obtained from ATCC (Manassas, VA, USA) and cultured in DMEM supplemented with 10% (v/v) FBS, 100 mg/Lpenicillin, and 100 mg/L streptomycin in 5% CO_2_ at 37 ℃. For transient transfection, approximately 2.5×10^6^ cells weremixed with 2 µg plasmids in 200 µL transfection buffer, and electroporation was carried out with a Scientz-2C electroporation apparatus (Scientz Biotech, Ningbo, China). The experiments were carried out 24 hours after transfection. HEK293 cell lines stably expressing Gα_16_ were constructed by our laboratory.

### Calcium Mobilization Assay

For OXGR1, plasmids were transfected into HEK293 (other receptors were transfected into HEK293 stably expressing Gα_16_) and seeded at a density of 4×10^4^ per well into 96-well culture plates and incubated for 24 h at 37 ℃ in 5% CO2. The next day, Cells were incubated with 2 μM/L Fluo-4 AM in HBSS supplemented with 5.6 mM/L D-glucose and 250 μM/L sulfinpyrazone at 37°C for 45 min. After washing, cells were added with 50 μL HBSS and incubated at room temperature for 10 min, then 25 μL agonist buffer was dispensed into the well using a FlexStation III microplate reader (Molecular Devices), and intracellular calcium change was recorded at an excitation wavelength of 485 nm and an emission wavelength of 525 nm. Three independent measurements were repeated, i.e. three different transfections. EC_50_ and E_max_ values for each curve were calculated by Prism 8.0 software (GraphPad Software).

### Surface expression analysis

After transfection, cells were seeded in 96-well plates. The next day, cells were washed with PBS, fixed with 4% PFA for 15 min, and then blocked with 2% BSA for 1 h. Next, cells were incubated with the polyclonal anti-HA (Sigma, H6908) overnight at 4°C and then horseradish peroxidase (HRP)-conjugated anti-rabbit antibody (CST, 7074S) for 1 h at room temperature. Then cells were washed and incubated with 50 μL tetramethylbenzidine (Sigma, T0440) for 30 min before the reaction was stopped with 25 μL TMB Substrate stop solution (Beyotime, P0215). Absorbance at 450 nm was quantified using a FlexStation III microplate reader (Molecular Devices).

### cAMP assay

In brief, cells expressing OXGR1 were seeded in plate at a density of 1×10^6^ cells/well and cultured overnight. After 24 h culture, cells were harvested and re-suspended in DMEM containing 500 µM IBMX at a density of 2×10^5^ cells/mL. Then cells were plated onto 384-well assay plates at a density of 1000 cells/5 µL/well. Another 5 µL buffer containing compounds at various concentrations were added to the cells and the incubation lasted for 30 min at 37 °C. Intracellular cAMP level was tested by a LANCE Ultra cAMP kit (PerkinElmer, TRF0264) and EnVision multiplate reader according to the manufacturer’s instructions. Three independent measurements were repeated, i.e. three different transfections.

### Expression and purification of Nb35

Nanobody-35 (Nb35) with a C-terminal 6×His tag, was expressed and purified^50^. Nb35 was purified by nickel affinity chromatography, followed by size-exclusion chromatography using a HiLoad 16/600 Superdex 75 column (Cytiva) and finally spin concentrated to 2 mg/mL.

### OXGR1 complexes expression and purification

The OXGR1, Gα_q_, Gβ_1_, Gγ_2_ and scFv16 were co-expressed in Trichoplusiani (Hi5) insect cells. Baculoviruses were prepared using the Bac-to-Bac Expression System (Invitrogen). Cells were infected with viruses at density of 3.0×10^6^ cell/mL with virus preparations for OXGR1, Gα_q_, Gβ_1_, Gγ_2_ and scFv16, at the ratio of 1:1:1:1. The infected cells were cultured at 27 °C for 48 h before collection by centrifugation and the cell pellets were stored at -80 °C until use. For the purification of the four OXGR1-G_q_ complexes, cell pellets were thawed in 20 mM HEPES pH 7.4, 100 mM NaCl, 5 mM MgCl_2_, 5 mM CaCl_2_, 25 mU/ml apyrase (Sigma-Aldrich), and protease inhibitor cocktail (TargetMol, USA). The suspensions were incubated overnight at 4 °C, in the presence of 100 mM AKG (TargetMol, USA) or 100 mM ITA (TargetMol, USA) or 100 mM SUC (TargetMol, USA) or 100 mM MA (TargetMol, USA), along with 10 μg/mL Nb35. Subsequently, 0.5% (w/v) n-dodecyl-β-d-maltoside (DDM, Anatrace) and 0.1% (w/v) cholesteryl hemisuccinate (CHS, Anatrace) were added to solubilize complexes for 2.5 h at 4°C. The supernatant was isolated by centrifugation at 30,000 r.p.m. for 30 min and then incubated overnight at 4 °C with pre-equilibrated TALON IMAC resin (Clontech). After batch binding, the TALON IMAC resin with immobilized protein complex was manually loaded onto a gravity flow column. The TALON IMAC resin was washed with 20 column volumes of 20 mM HEPES, pH 7.4, 100 mM NaCl, 5 mM CaCl_2_, 5 mM MgCl_2_, 30 mM imidazole, 10% glycerol, 0.1% LMNG (w/v), 0.02% CHS (w/v), 50 mM AKG or 50 mM ITA or 50 mM SUC or 50 mM MA and eluted with the same buffer plus 300 mM imidazole, 50 mM AKG or 50 mM ITAor 50 mM SUC or 50 mM MA. The mixture was then purified by SEC using a Superose 6 10/300 GL column (GE healthcare) in 20 mM HEPES, pH 7.4, 100 mM NaCl, 0.00075% (w/v) LMNG, 0.00025% (w/v) CHS and 50 mM AKG or 50 mM ITA or 50 mM SUC or 50 mM MA. The fractions of monomeric complex were collected and concentrated to 6.8-8.5 mg/ml for electron microscopy experiments.

### Cryo-EM grid preparation and data collection

For the preparation of cryo-EM grids, 3 μL of the purified protein were applied onto a glow-discharged holey Nitinol grid (M01 Au300-r1.2/1.3, Nanodim). Grids were plunge-frozen in liquid ethane using Vitrobot Mark IV (Thermo Fischer Scientific). Frozen grids were transferred to liquid nitrogen and stored for data acquisition. Cryo-EM imaging of the complex was performed on a Titan Krios at 300 kV in the Advanced Center for Electron Microscopy at Shanghai Institute of Materia Medica, Chinese Academy of Sciences (Shanghai China). A total of 6,034/5,264/5,096 movies for the AKG/ITA/MA-OXGR1-G_q_ complexes were collected on K3 direct electron detection device. Images were taken with a pixel size of 0.82 Å, a defocus ranging from -1.0 to -2.0 μm, and total dose of 50 e^−^ Å^2^ s^−1^, using the EPU software (FEI Eindhoven, Netherlands). For the SUC-OXGR1-G_q_ complexes, 8,999 movies were collected on Falcon 4 direct electron detection device. Images were taken with a pixel size of 0.73 Å, a defocus ranging from -1.0 to -2.0 μm, and total dose of 50 e^−^ Å^2^ s^−1^ over 2.5 s exposure on each EER format movie, using the EPU software (FEI Eindhoven, Netherlands). Each movie was divided into 36 frames during motion correction.

### Image processing and map construction

The single particle analysis of OXGR1-G_q_ complex was performed with cryoSPARC v4^51^. Dose-fractionated imagestacks were subjected to motion correction by MotionCor2 ^52^. Contrast transfer function (CTF) parameters for micrograph were estimated by patch CTF estimation. For the AKG-OXGR1-G_q_ complex, auto-picked particles were extracted from a binned dataset with a pixel size of 1.64 Å and subjected to reference-free 2D classification to remove poorly defined particles. After several rounds of 2D classification, a total of 2,058,837 particles were obtained. These particles were then projected with hetero-refinement, and two subsets of well-defined particles, totally 1,609,345 particles, were re-extracted at a pixel size of 0.82 Å. Considering the flexibility between the receptor and G proteins, we generated a mask focus on the receptor, and used as focused mask in following steps. The extracted particles were subjected to 3D classification with focused mask. Two subset consisting of 405,687 particles, showing the best performance on the ligand, was selected for further non-uniform refinement and local refinement. The final density map was obtained with an overall resolution of 2.66 Å. Additional local refinement focus on the receptor produced a local map with a resolution of 2.89 Å.

For the ITA-OXGR1-G_q_ complex, a total of 1,906,730 particles were picked from template and Topaz extracted particles through several rounds of 2D classification. These particles were then projected with hetero-refinement, one subset with well-defined particles were re-extracted at a pixel size of 0.82 Å. A total of 660,367 particles were subjected to 3D classification with focused mask on the receptor. The three subsets consisting of 405,209 particles, showing the best performance on the ligand, was selected for further non-uniform refinement and local refinement. The final density map was obtained with an overall resolution of 2. 65 Å. Additional local refinement focus on the receptor produced a local map with a resolution of 2.90 Å. For the MA-OXGR1-G_q_ complex, a total of 1,945,913 particles were picked from template and Topaz extracted particles through several rounds of 2D classification. These particles were then projected with hetero-refinement, one subset with well-defined particles were re-extracted at a pixel size of 0.82 Å. A total of 1,083,657 particles were subjected to 3D classification with focused mask on the receptor. The subset consisting of 436,647 particles, showing the best performance on the ligand map, was selected for further non-uniform refinement and local refinement. The final density map was obtained with an overall resolution of 2. 73 Å. Additional local refinement focuses on the receptor produced a local map with a resolution of 2.97 Å. For the SUC-OXGR1-G_q_ complex, a total of 3,890,029 particles were picked from template and Topaz extracted particles through several rounds of 2D classification. These particles were then projected with hetero-refinement, two subsets with well-defined particles were re-extracted at a pixel size of 0.73 Å. A total of 2,250,721 particles were subjected to 3D classification. The subset consisting of 360,330 particles, showing the best performance on the ligand map, was selected for further non-uniform refinement and local refinement. The final density map was obtained with an overall resolution of 2.64 Å. Additional local refinement focus on the receptor produced a local map with a resolution of 2.70 Å. All resolutions are determined by a Fourier shell correlation of 0.143.

### Model building and refinement

For all OXGR1-G_q_ complexes, the AlphaFold2 structure of OXGR1^53^, were used as the initial model for model rebuilding, and refinement against the EM mapsthat focus on theligand bound receptor. And the structure of G_q_ protein (PDB code: 8XOF) were used as the initial model for model rebuilding and refinement against the EM maps of full map of ligand bound OXGR1-G_q_ complex.

The model was docked into the electron microscopy density map using Chimera^54^, followed by iterative manual adjustment and rebuilding in COOT^55^ and ISOLDE^56^. Real space and reciprocal space refinements were performed using Phenix programs^57^. The model statistics were validated using MolProbity^58^. The final refinement statistics were validated using the module “comprehensive validation (cryo-EM)” in Phenix. The final refinement statistics are provided in Supplementary Table 1. Structural figures were prepared in ChimeraX^59^ and PyMOL (https://pymol.org/2/).

### MD simulations

The simulation systems were constructed based on the OXGR-ITA, OXGR-AKG, OXGR-SC, and OXGR-MA complexes. The F130D mutation was introduced using PyMOL, and G proteins were excluded from the systems. System information was summarized in Supplementary Table 5. Each complex was embedded into a 75 × 75 Å POPC lipid bilayer using the packmol-memgen software and solvated with a 12 Å layer of water^60^. The ionic strength was maintained at 0.15 mol/L NaCl, with counterions added to neutralize the system. The FF19SB, Lipid21, and GAFF2 force fields were applied to model amino acids, lipids, and ligands, respectively^61–63^. Such a system composition has been confirmed in multiple studies ^64–66^. Each system underwent energy minimization, followed by heating and equilibration using standard protocols^64,67^. In the heating phase, the system starts from an initial temperature of 0 K and is gradually warmed to 300 K over 150,000 steps (300 ps with a 2 fs timestep) using the NVT ensemble (constant volume and temperature). The Langevin thermostat controls the temperature, and positional restraints are applied to keep the system fixed with a force constant of 10.0 kcal/mol/Å². For the equilibration phase, the system continues in the NVT ensemble at 300 K for 350,000 steps (700 ps with a 2 fs timestep). The Langevin thermostat maintains the temperature, and the same positional restraints are applied to the system. This step allows the system to stabilize at the target temperature before the production run. Although transitioning to the NPT ensemble during equilibration is a common practice to allow for volume and density adjustment, we retained NVT to maintain strict control over the system’s initial relaxation under positional restraints, given the prior minimization step.

Six independent production runs of 1000 ns each were performed using pmemd.cuda in Amber22 under the NVT ensemble at 300 K and 1 atm^68^. Long-range electrostatic interactions were calculated using the Particle Mesh Ewald (PME) method, while short-range electrostatic and van der Waals interactions were truncated at a 10 Å cutoff. The SHAKE algorithm and hydrogen mass repartitioning were employed to constrain bonds involving hydrogen atoms, enabling a 4 fs timestep. Distance calculationswere conducted using CPPTRAJ^69^. Binding free energies were estimated using the MMPBSA.py tool^70^. Reference structure for root mean squared deviation (RMSD) was the cryo-EM strcuture and the residue RMSD is calculated on heavy atoms after the whole structure was aliganed to all Cα coordinates. Binding free energies were estimated using the MMPBSA.py tool^70^. During free energy calculation, we did not include the entropy contribution to the binding free energy in order to reduce computational cost and avoid the known convergence challenges associated with normal mode or quasi-harmonic analyses. Consequently, these calculations reflect relative trends rather than absolute binding free energy values. The MMPBSA calculations were conducted using implicit solvent, specifically the Generalized Born model with igb=5, which corresponds to GB-OBC model II. This model uses optimized parameters and is recommended for use with the mbondi2 radii set. Such combination of hydrogen mass repartitioning and binding free energy calculation has been confirmed in^71^.

### Molecular docking

The AKG/ITA/SUC/MA-OXGR1 complex structures were used as the starting point for molecular docking. The OXGR1 was isolated from the G protein complex and prepared using the Protein Preparation Wizard in Schrödinger’s Maestro. The ligands, AKG/ITA/SUC/MA, were prepared using the LigPrep module, ensuring proper bond orders and protonation states. Hydrogen atoms were added to the protein, disulfide bonds were defined, and heteroatom states of residues were assigned using Epik at pH 7.0 ± 2.0. Residue protonation states were determined using PROPKA1^72^. Grid files were generated based on the ligand-binding pocket of the receptor. The ligands were then docked into the prepared grids using the standard precision mode of the Glide program. The docking pose with the highest docking score was selected as the final result for further analysis.

## Acknowledgements

This work was supported by grants from the National Natural Science Foundation of China (32130022 to H.E.X., 82121005 to X.X. and H.E.X., 82330113 to X.X., 82304579 to S.G.); The National Key R&D Program of China (2022YFC2703105 to H.E.X., 2022YFA1302900 to W.Y.); CAS Strategic Priority Research Program (XDB37030103 to H.E.X.); Shanghai Municipal Science and Technology Major Project (2019SHZDZX02 to H.E.X.); Shanghai Municipal Science and Technology Major Project (H.E.X.); the Lingang Laboratory, Grant No.LG-GG-202204-01 (H.E.X.); the Natural Science Foundation of Shanghai (25ZR1402552 to H.L.). The cryo-EM data were collected at the Shanghai Advanced Electron Microscope Center, Shanghai Institute of Material Medica. We thank Q. Yuan., K. Wu., W. Hu., S. Li. And S. Zhang. for providing technical support and assistance during data collection at the Shanghai Advanced Electron Microscope Center, Shanghai Institute of Material Medica.

## Author contributions

H.E.X. and H.L. initiated the project. X.Z., S.G., Y.W., W.Y. H.L., X.X., and H.E.X. administrated the project. X.Z. screened the expression constructs and optimized the purification conditions of protein complexes, prepared protein samples for cryo-EM grid preparation, and participated in figure preparation. Y.L., Y.T. and S.G. performed the cell-based functional assays and participated in figure preparation. H.L. performed the cryo-EM data calculations, model building. X.H. performed the MD simulations and participated in figure preparation. C.L., Y.W., Y.G. and J.Y. participated in the functional studies and protein purification. Q.Y. participated in EM data calculations. W.H. and K.W. screened the EM sample and collected EM movies. H.L. and X.Z. prepared the figures and wrote the draft. Y.W., W.Y. revised the manuscript. H.E.X., H.L. and X.X. conceived and supervised the project and wrote the manuscript with inputs from the authors.

## Data availability

Density maps and structure coordinates have been deposited in the Electron Microscopy Data Bank (EMDB) and the Protein Data Bank (PDB) with accession codes EMD-63580 and 9M1R for AKG-OXGR1-G_q_ complexand EMD-64590 and 9UXQfor local refinement of AKG bound OXGR1; EMD-63581 and 9M1S for ITA-OXGR1-G_q_ complex and EMD-64589 and 9UXP for Local refinement of ITA bound OXGR1; EMD-63583 and 9M1U for SUC-OXGR1-G_q_ complex and EMD-64587 and 9UXN for local refinement of Succinate bound OXGR1; EMD-63582 and 9M1T for MA-OXGR1-G_q_ complex and EMD-64588 and 9UXO for local refinement of maleic acid bound OXGR1; Source data is provided with this paper. Simulation trajectories and inputs can be downloaded from https://doi.org/10.6084/m9.figshare.28946102.v2.

