## Extended data figure for "Molecular architecture of OXGR1 reveals evolutionary conserved mechanisms for metabolite surveillance"

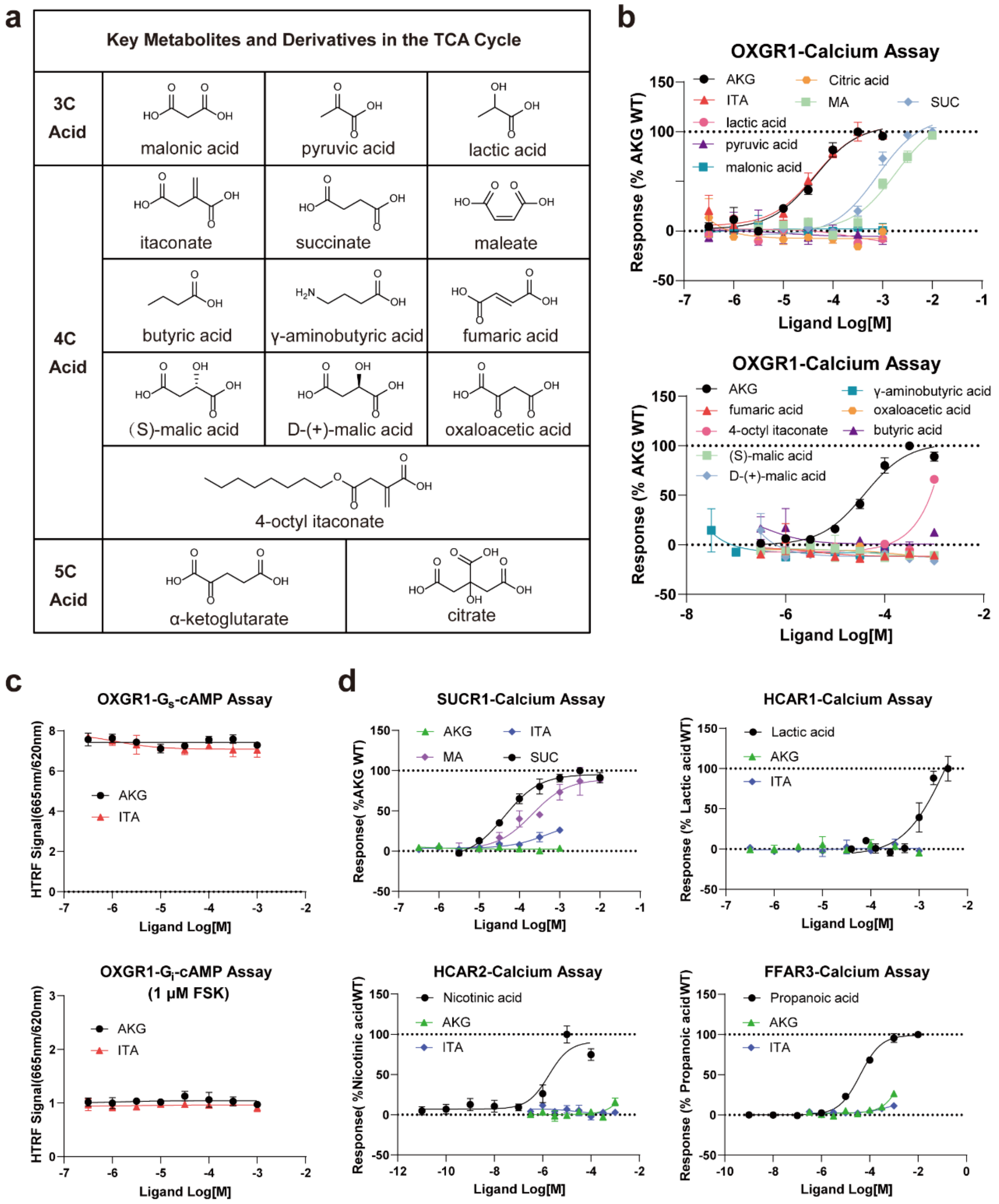


**Extended Data Fig. 1| Functional characterization of TCA metabolites and derivatives in the activation of OXGR1.**

**(a)** Chemical structures of investigated metabolites and derivatives;

**(b-d)** Concentration-response curves of the 15 metabolites and derivatives in activating OXGR1 through the calcium assay **(b)**, assessment of AKG and ITA activating OXGR1 through $G_i$/$G_s$ pathway in cAMP assay **(c)**, and Concentration-response curves of the ITA and AKG in activating SUCR1, HCAR1/HCAR2, FFAR3 **(d)**. Data are mean ± S.E.M. from three independent experiments ($n = 3$).

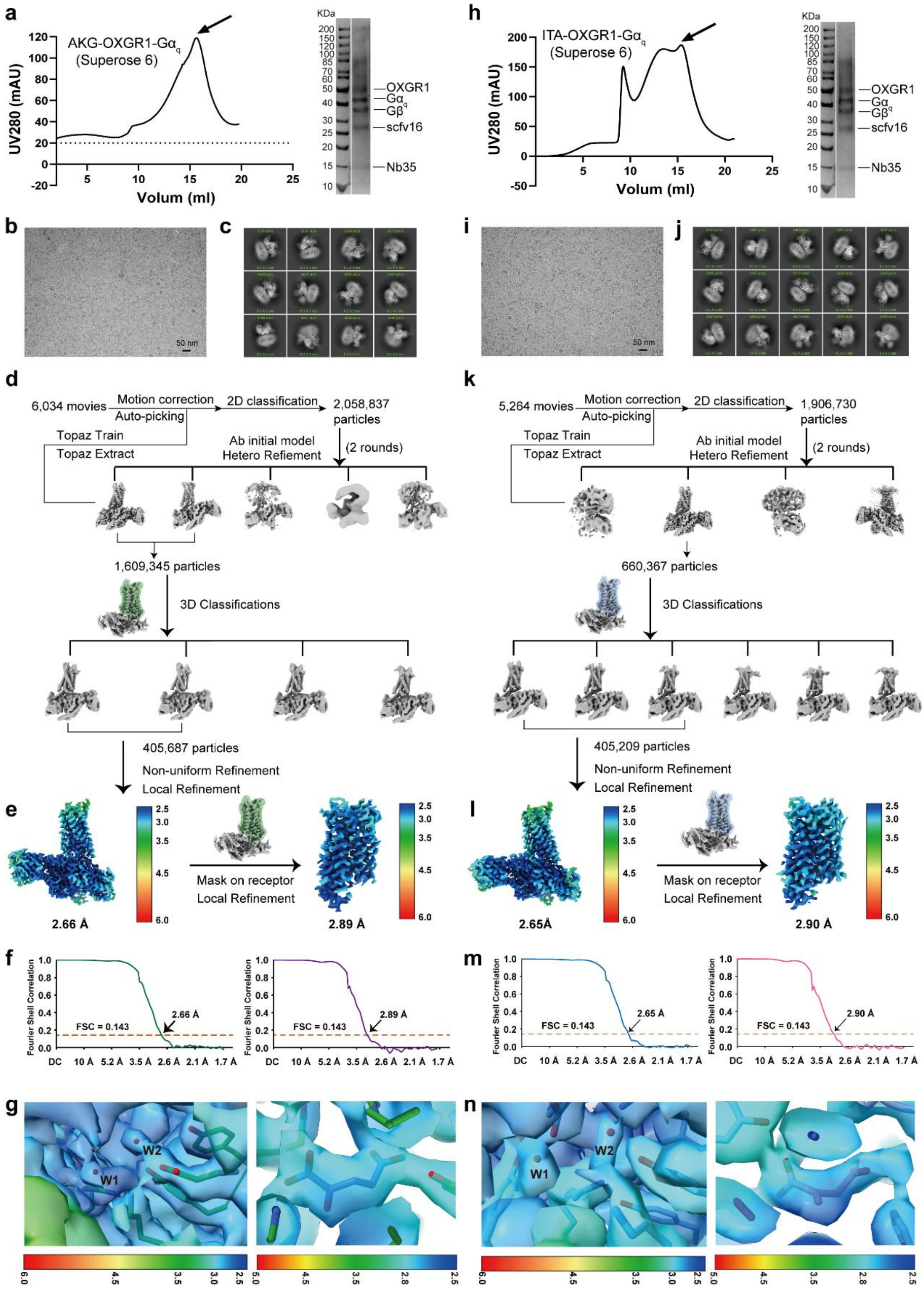


**Extended Data Fig. 2| Purification and data-procession of AKG-OXGR1-$G_q$ and ITA-OXGR1-$G_q$ complex.**

**(a, h)** Representative size exclusion chromatography (SEC) profiles and SDS-PAGE analysis of OXGR1-$G_q$ complex activated by AKG **(a)** and ITA **(h)**. Experiment was repeated at least three times with similar results.

**(b-c, and i-j)** Representative cryo-EM image and 2D classification averages of AKG-OXGR1-$G_q$ complex

**(d, and k)** Cryo-EM data processing flowcharts of AKG-OXGR1-$G_q$ **(d)** and ITA-OXGR1-$G_q$ complexes **(k)**.

**(e, and l)** The global and local receptors density maps of AKG-OXGR1-$G_q$ **(e)** and ITA-OXGR1-$G_q$ **(l)** colored by local resolutions.

**(f, and m)** The Fourier shell correlation (FSC) curves of AKG-OXGR1-$G_q$, AKG-OXGR1 **(f)** and ITA-OXGR1-$G_q$, ITA -OXGR1 **(m)**. The global resolution of the final processed density map estimated at the FSC = 0.143 is 2.66 Å and 2.65 Å. The local resolution of the final processed receptor density map estimated at the FSC = 0.143 is 2.89 Å and 2.90 Å.

**(g, and n)** The density of water molecules (W1/W2) and AKG **(g)**/ITA **(n)** in the in the OXGR1 binding pocket, colored by local resolution, are shown at a contour level of 0.3.

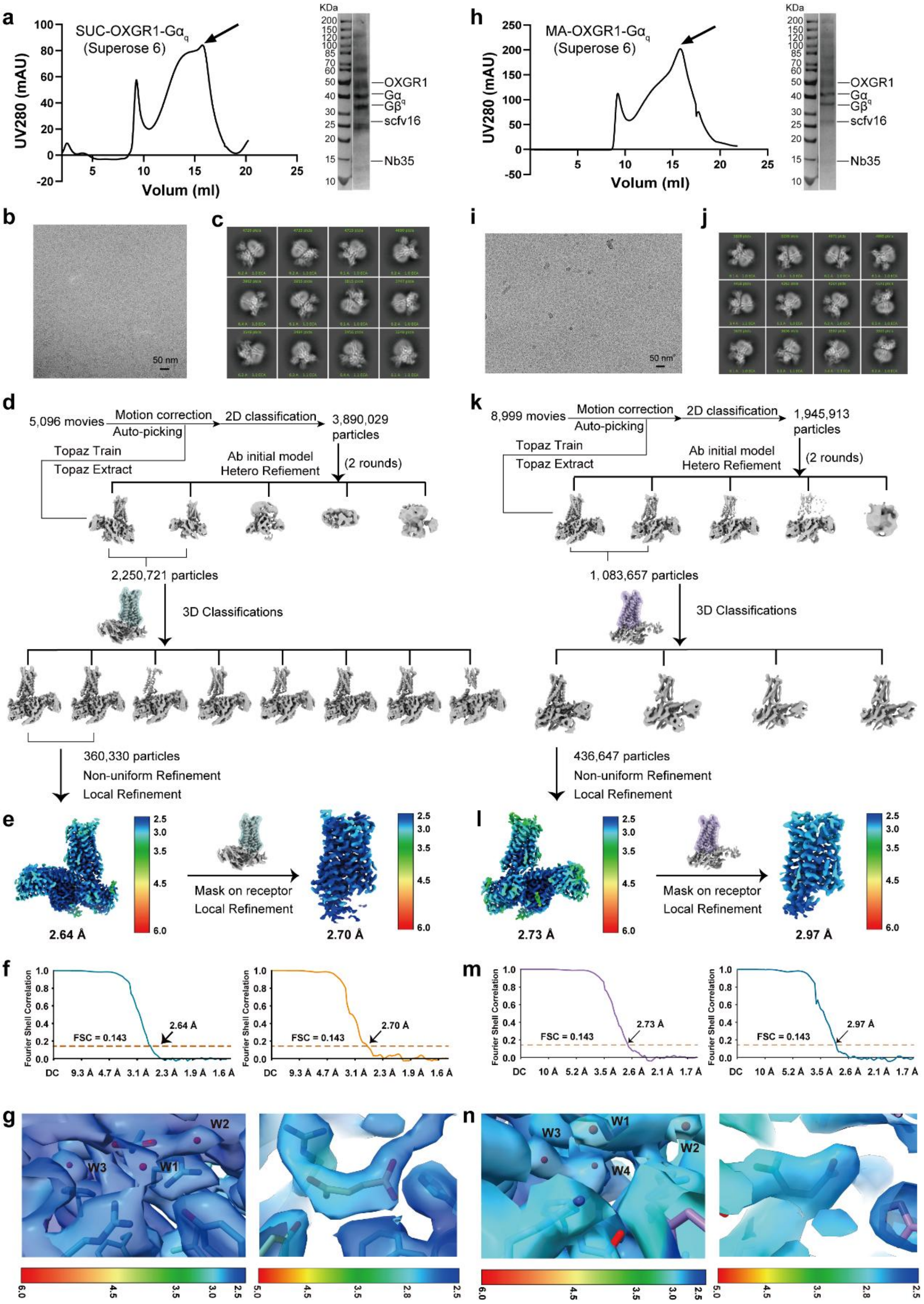


**Extended Data Fig. 3| Purification and data-procession of SUC-OXGR1-$G_q$ and MA-OXGR1-$G_q$ complex.**

**(a, h)** Representative size exclusion chromatography (SEC) profiles and SDS-PAGE analysis of OXGR1-$G_q$ complex activated by SUC **(a)** and MA **(h)**. Experiment was repeated at least three times with similar results.

**(b, and i)** Representative cryo-EM image of SUC-OXGR1-$G_q$ **(b)** and MA-OXGR1-$G_q$ complex **(i)**.

**(c, and j)** Representative 2D classification averages of SUC-OXGR1-$G_q$ **(c)** and MA-OXGR1-$G_q$ complex **(j)**.

**(d, and k)** Cryo-EM data processing flowcharts of SUC-OXGR1-$G_q$ **(d)** and MA-OXGR1-$G_q$ complexes **(k)**.

**(e, and l)** The global density map of SUC-OXGR1-$G_q$ **(e)** and MA-OXGR1-$G_q$ **(l)** colored by local resolutions.

**(f, and m)** The Fourier shell correlation (FSC) curves of SUC-OXGR1-$G_q$ **(f)** and MA-OXGR1-$G_q$ **(m)**. The global resolution of the final processed density map estimated at the FSC = 0.143 is 2.64 Å and 2.73 Å. The local resolution of the final processed receptor density map estimated at the FSC = 0.143 is 2.70 Å and 2.97 Å.

**(g, and n)** The density of water molecules (W1/W2/W3/W4) and SUC **(g)**/MA **(n)** in the in the OXGR1 binding pocket, colored by local resolution, are shown at a contour level of 0.3.

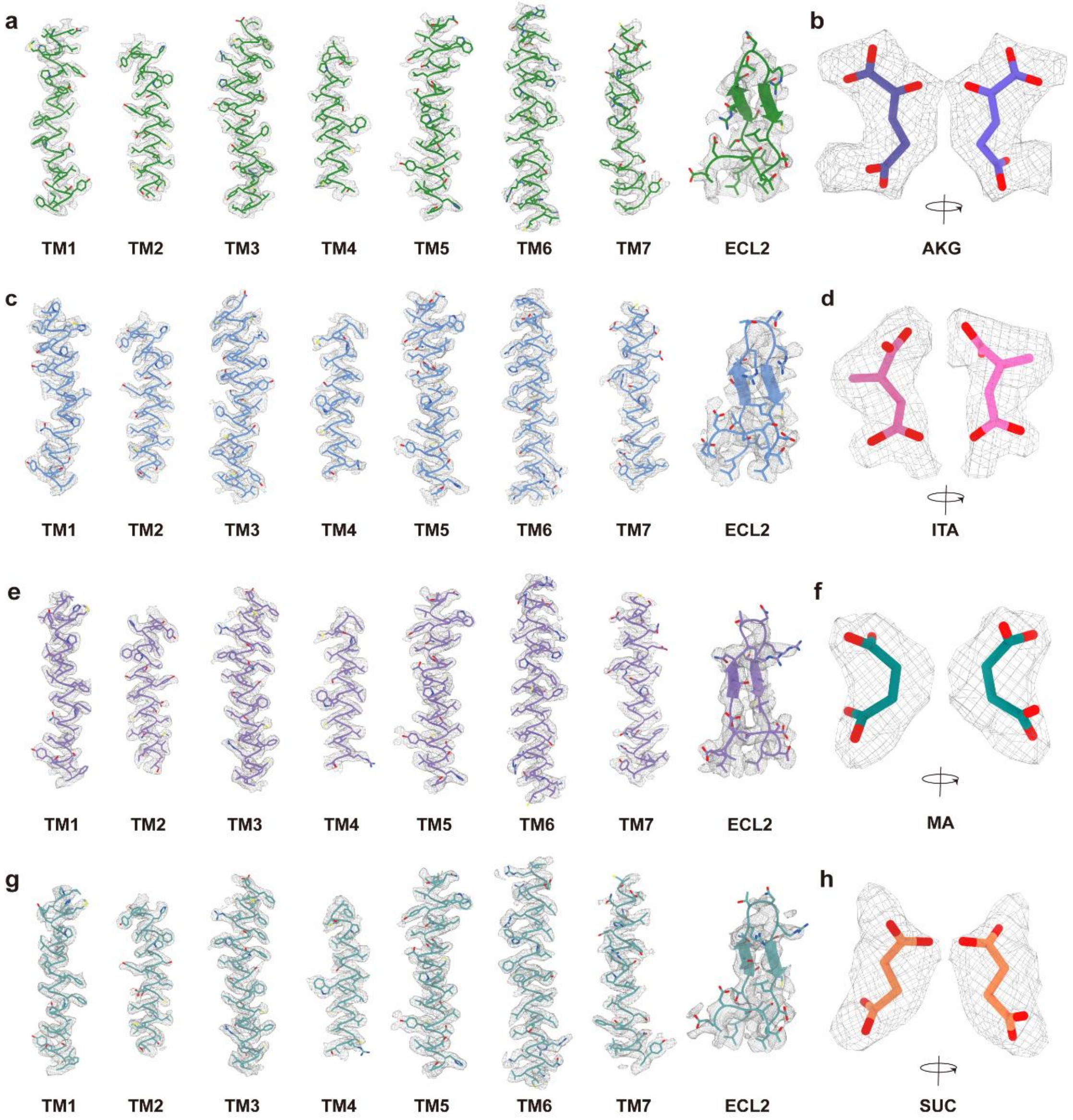


**Extended Data Fig. 4| The density maps of protein and four ligands in OXGR1-$G_q$ complex.**

**(a-h)** The density maps of helices TM1-TM7 of transmembrane domain, extracellular loop ECL2 of OXGR1**(a, c, e, g)** in OXGR1-$G_q$ complex in the presence of AKG **(b)**/ITA**(d)** /SUC**(f)** /MA**(h)**. All protein density maps are shown at contour level of 0.5. The densities of respective ligands have been extracted from their local structures of receptors and shown in mesh presentation. All ligand density maps are shown at contour level of 0.3, and the local resolutions of AKG/ITA/SUC/MA-bound OXGR1 are 2.89Å, 2.90 Å, 2.70 Å, 2.97 Å, respectively.

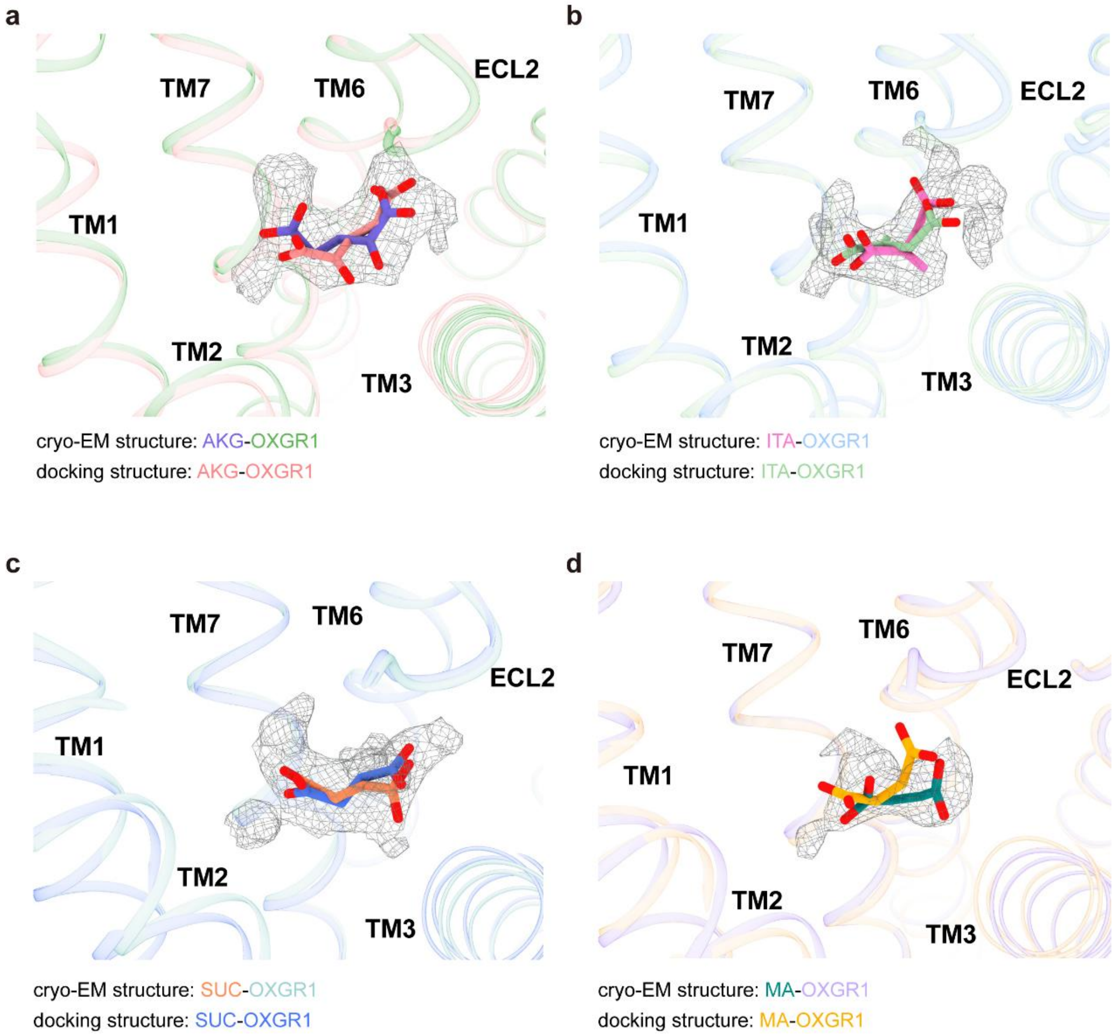


**Extended Data Fig. 5| Structural validation of four ligand-binding modes in OXGR1 by molecular docking and cryo-EM density maps.**

**(a-b)** Comparison of AKG **(a)**, ITA **(b)**, SUC **(c)**, and MA **(d)** binding postures in molecular docking and structural modeling of OXGR1. Each ligand is aligned with the corresponding cryo-EM density represented in mesh, validating the proposed binding conformations. The densities of respective ligands have been extracted from their local structures of receptors and shown in surface presentation. All ligand density maps are shown at contour level of 0.3, and the local resolutions of AKG/ITA/SUC/MA-bound OXGR1 are 2.89 Å, 2.90 Å, 2.70 Å, 2.97 Å, respectively.

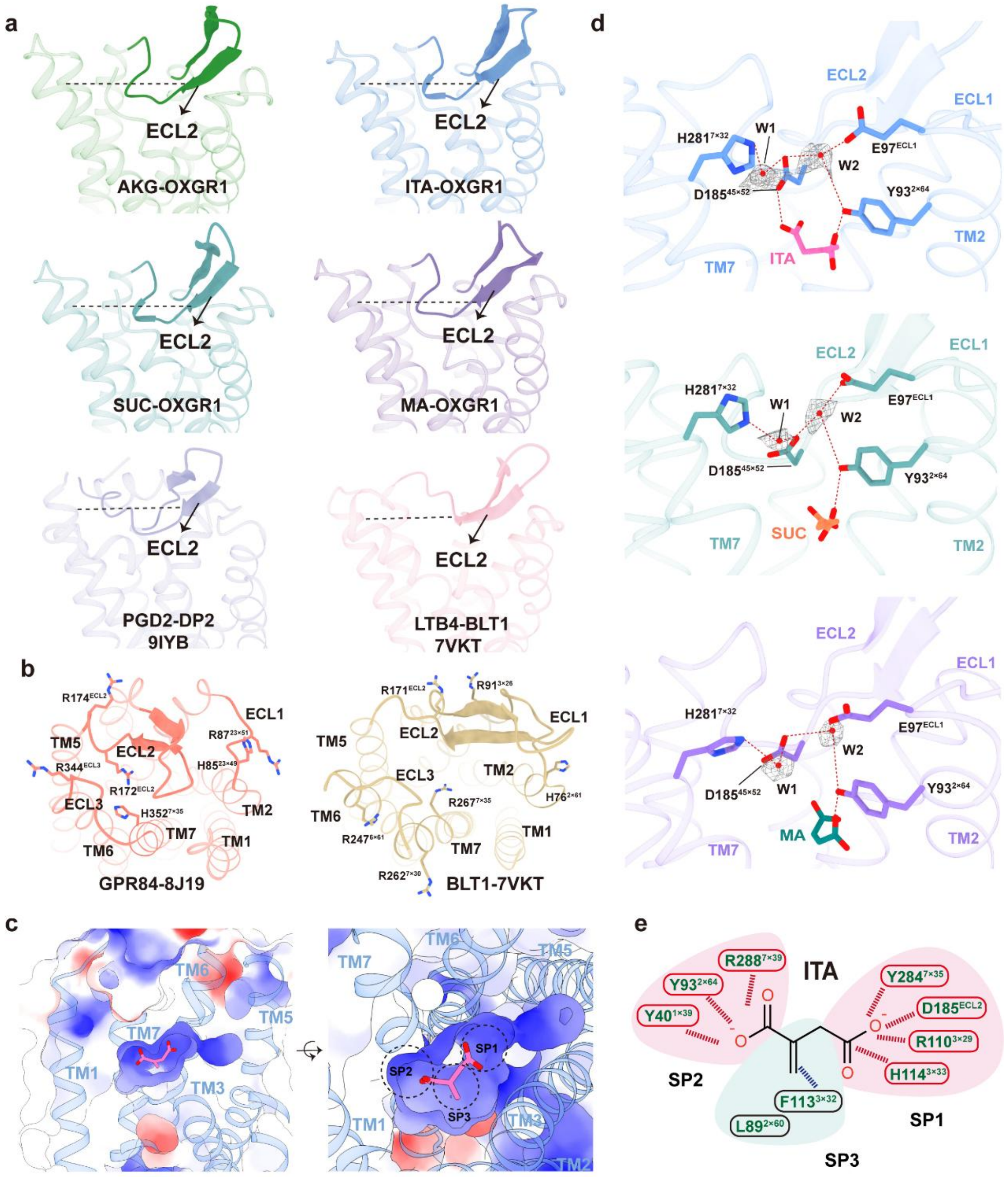


**Extended Data Fig. 6| Structural features of OXGR1-$G_q$ complex.**

**(a)** Similar structure feature of ECL2-β-hairpin partially buried within the orthosteric pocket.

**(b)** The structures of BLT1 and GPR84 reveal a convergent “cationic lure” ligand entry mechanism featuring, as observed in the case of OXGR1 in Fig. 2e-f.

**(c)** The ligand binding pocket, with ITA-OXGR1 as an additional description, is characterized by a polar and positively charged nature.

**(d)** Additional polar hydrogen-bond network in ITA/SUC/MA-OXGR1 structures mediated by two water molecules, W1 and W2, is surrounded by key residues shown in sticks. The ordered water molecules (W1/W2) are overlaid with their corresponding cryo-EM density represented in mesh, and their density maps are shown at contour level of 0.35. Key polar interactions are shown with red dashed lines.

**(e)** Schematic illustration of the ligand recognition mechanism of OXGR1 using ITA as a supplementary example. The carboxy groups of ITA are deprotonated. Key polar interactions in the pink region are shown with red dashed lines and red circles, and hydrophobic interactions in the cyan region are shown with blue dashed lines and black circles.

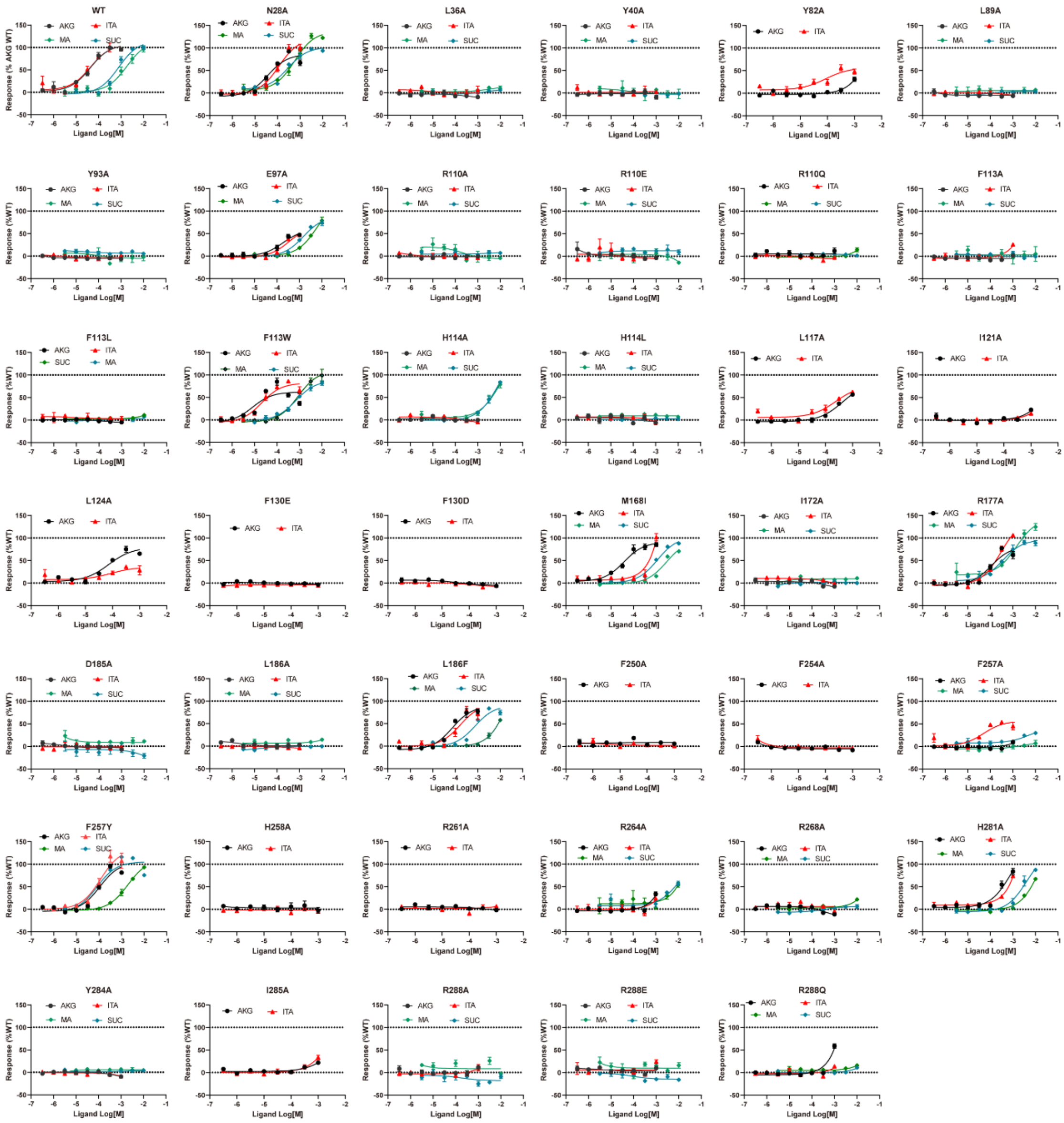


**Extended Data Fig. 7| Mutation and functional assay indicates ligand effects on the activation of mutated OXGR1 variants.**

Concentration-response curves show ligand-induced activation of mutated OXGR1 variants corresponding to ligand binding. Data are mean ± S.E.M. from three independent experiments (*n* = 3). Three independent measurements refer to three different transfections.

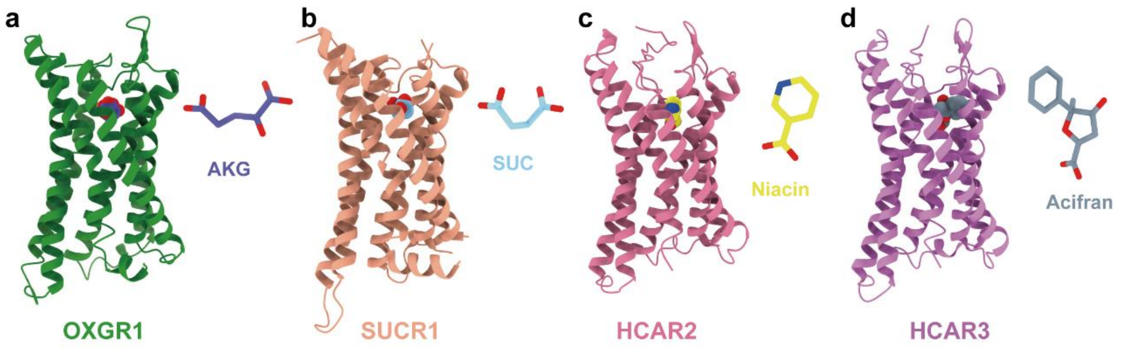


**Extended Data Fig. 8|** Different recognition modes of ligands in carboxylic acid-sensing receptors.

Binding pose of ligands in cryo-EM structures of OXGR1-AKG (**a**), SUC-SUCR1 (**b,** PDB:8YKW), Niacin-HCA2 (**c**, PDB:8K5B), and Acifran-HCAR3 (**d**, PDB:8IHK). OXGR1 and SUCR1 share a horizontal ligand-binding mode (**a** and **b**), in contrast to the vertical binding pose adopted by HCAR2 and HCAR3(**c** and **d**).

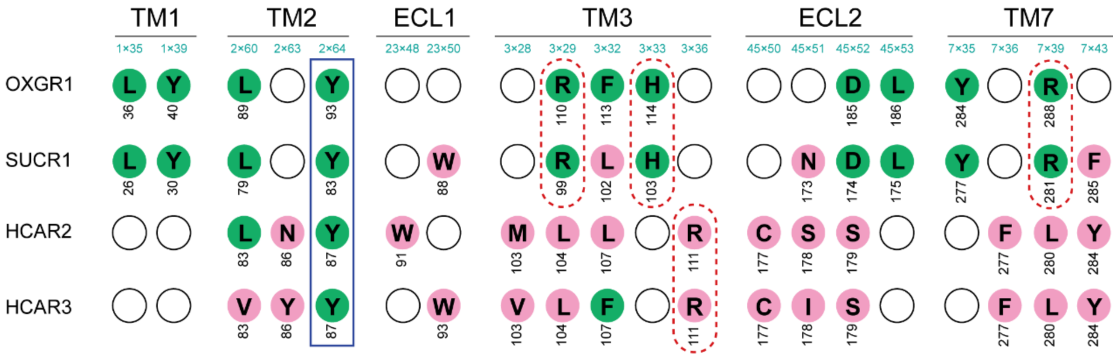


**Extended Data Fig. 9| Key basic residues in OXGR1 binding pocket underlies selective dicarboxylate recognition.**

Sequence alignment of OXGR1, SUCR1, HCAR2, and HCAR3 binding residues reveals a basic residue-rich environment in OXGR1. This feature is shared with SUCR1 but absent in HCAR2/3, which is consistent with ligand orientation in Fig. 5a-c. Identical residues to the OXGR1 binding pocket are highlighted with green backgrounds; divergent residues with pink. Fully conserved residues across the receptor family are blue-boxed; non-conserved basic residues are red-dashed-circled.

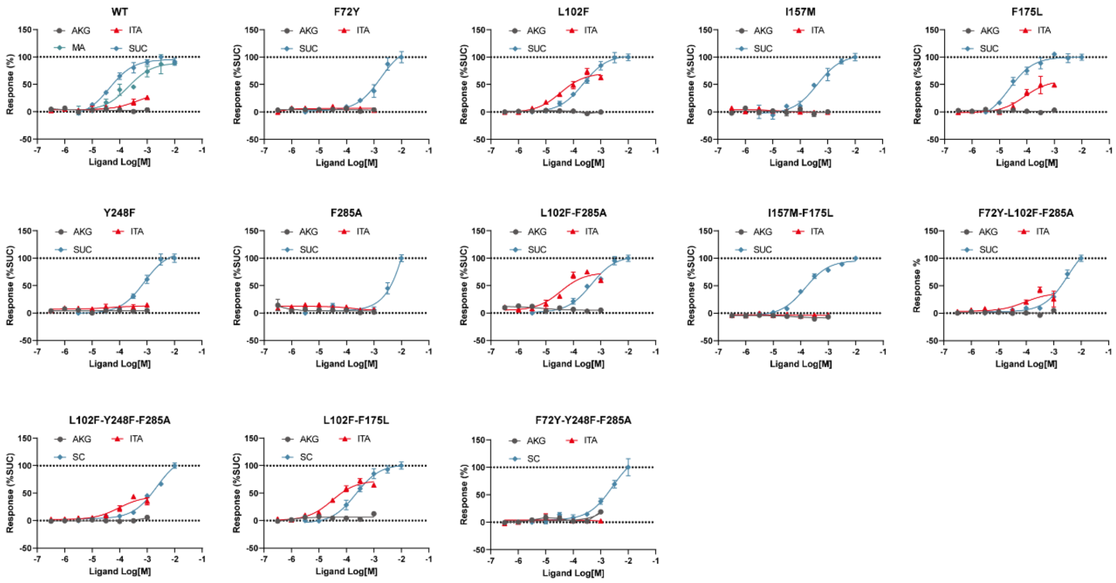


**Extended Data Fig. 10| Mutation and functional assay indicates ligand effects on the activation of mutated SUCR1 variants.**

Concentration-response curves show ligand-induced activation of mutated SUCR1 variants corresponding to ligand binding. Data are mean ± S.E.M. from three independent experiments (*n* = 3). Three independent measurements refer to three different transfections.

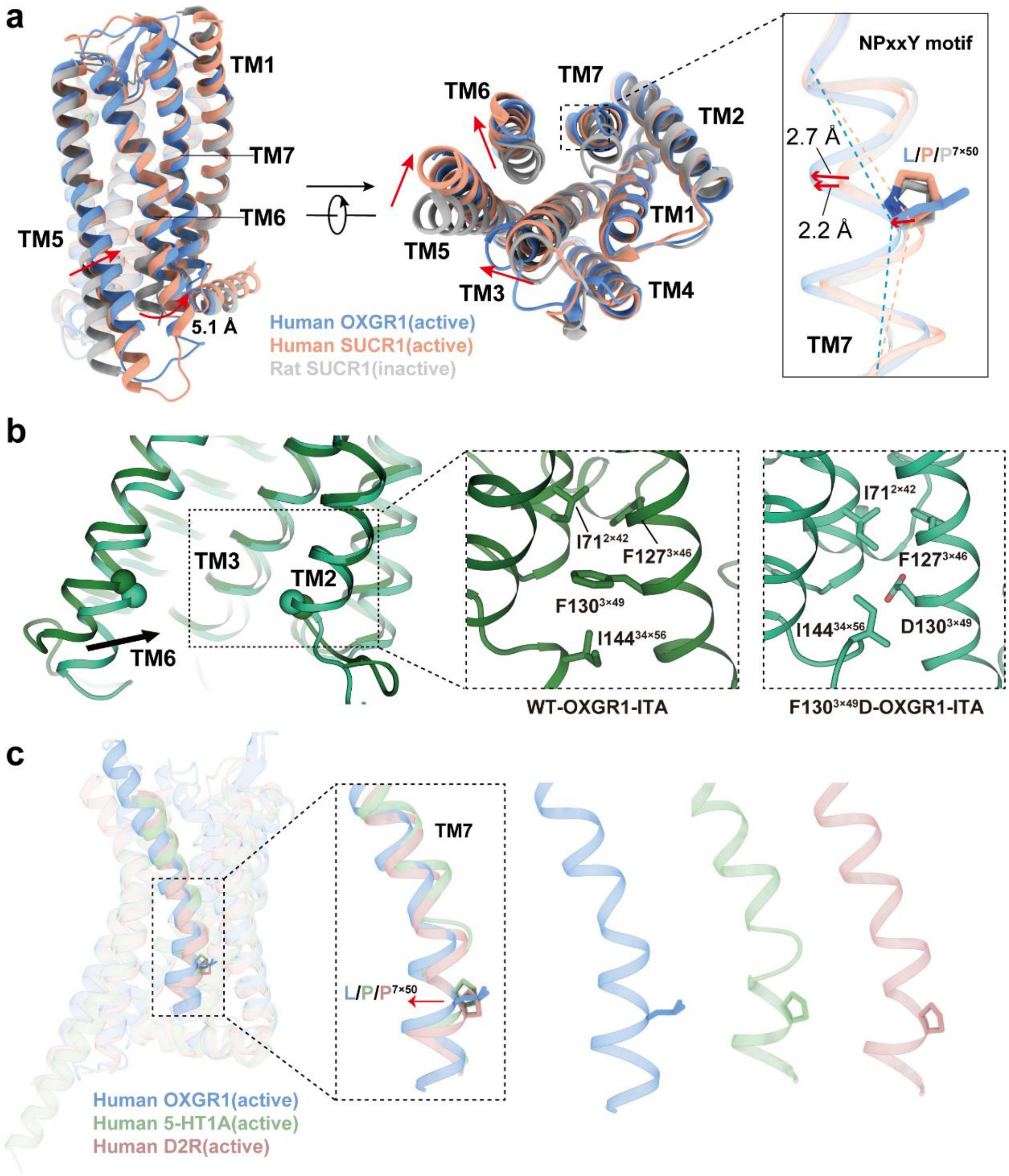


**Extended Data Fig. 11| Unique activation motifs contribute to OXGR1 signaling.**

**(a)** Distinct conformational changes in OXGR1 activation. Structural comparison of the ITA-bound OXGR1 with the inactive and active SUCR1 revealed a pronounced outward displacement of the cytoplasmic end of TM6, a rotation of TM5, as well as the inward movement of TM7, all hallmark features of receptor activation.

**(b)** FRY motif drives hydrophobic pocket remodeling and outward movement of TM6. Inward movement of TM6 in F130$^{3x49}$D system is observed in comparison with WT system, and the zoom-in view of hydrophobic pocket around F/D$^{3x49}$ reveals that OXGR1's FRY motif stabilizes the active state through enhanced hydrophobic packing.

The present findings indicate that the FRY motif of OXGR1 facilitates its activation.

**(c)** Structural comparison of OXGR1 with 5-$HT_{1A}$ and D2R highlights OXGR1's divergent rearrangement in the NPxxY motif, which differs from the canonical Class A GPCR mechanisms.

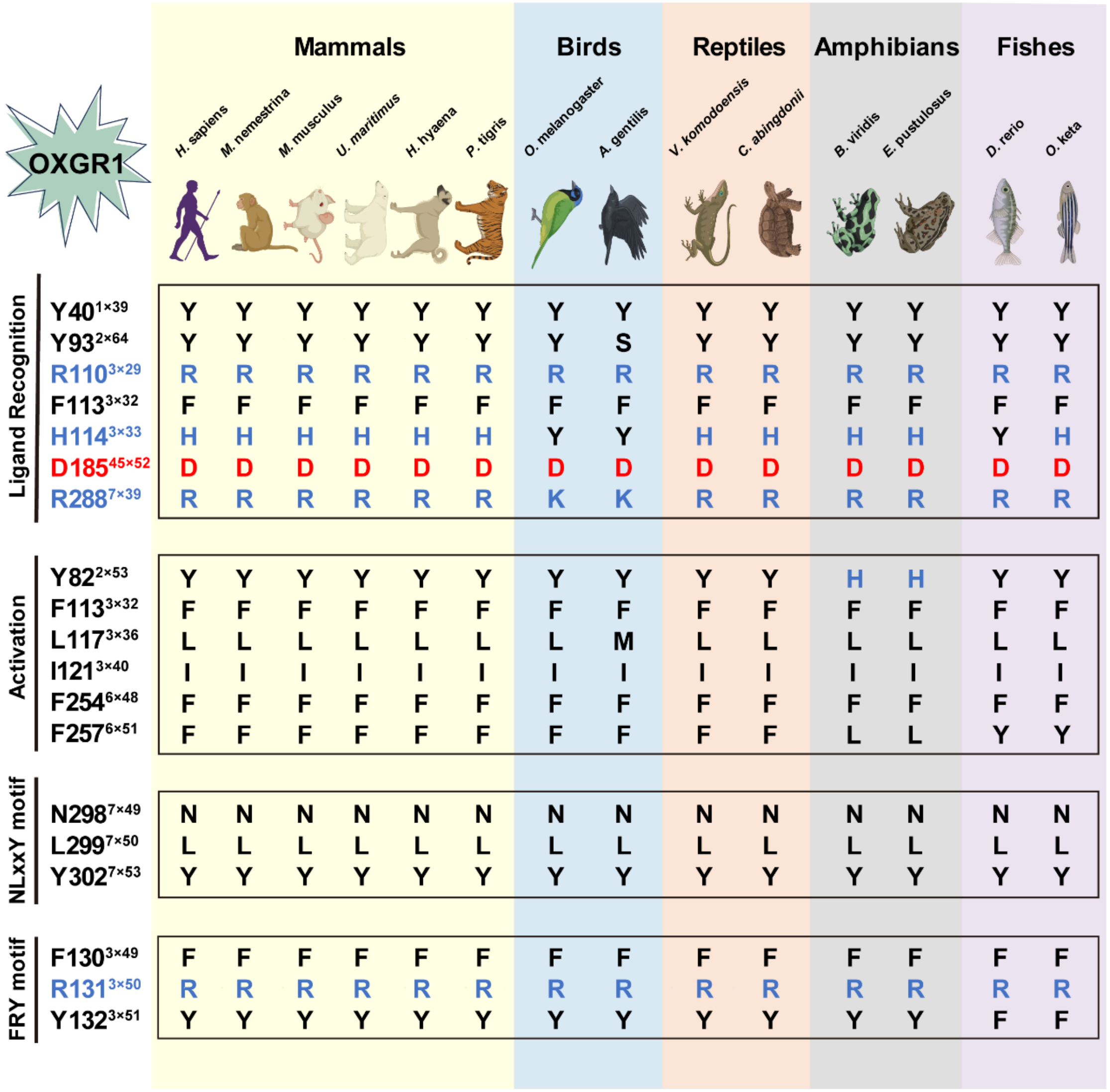


**Extended Data Fig. 12| Evolutionary conservation of OXGR1 function in vertebrate.**

Sequence alignment of key residues involved in the ligand recognition, activation mechanism, $F^{3x49}$-$R^{3x50}$-$Y^{3x51}$ motif (D/ERY motif in common GPCRs) and N-$L^{7.50}$-xx-$Y^{7.53}$ motifs (NPxxY motif in common GPCRs) of OXGR1 among different species in vertebrates. Highlight basic amino acids in blue, acidic amino acid in red, and other residues in black. The animal images were generated using BioRender (https://biorender.com).

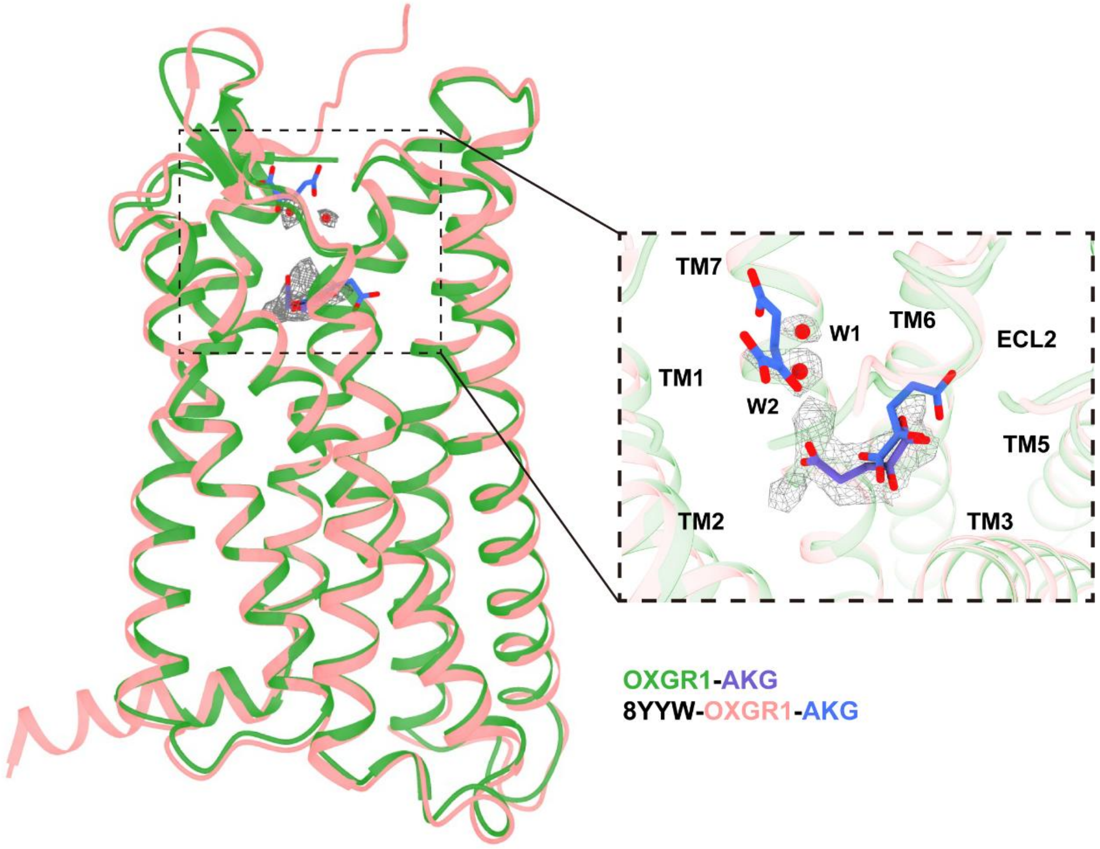


**Extended Data Fig. 13|** Comparison of ligand recognition between OXGR1-AKG complex and recently released OXGR1-AKG structure (PDB ID: 8YYW).

The AKG of our cryo-EM structure overlaid on corresponding density map in gray mesh shown at contour level of 0.3, with AKG in the orthosteric pocket. Superposition with 8YYW showing partial overlap of primary AKG. The density near ECL2 resolves as two water molecules in our 2.89 Å cryo-EM structure.
